# Partial endoreplication stimulates diversification in the species-richest lineage of orchids

**DOI:** 10.1101/2020.05.12.091074

**Authors:** Zuzana Chumová, Eliška Záveská, Jan Ponert, Philipp-André Schmidt, Pavel Trávníček

**Affiliations:** Czech Academy of Sciences, Institute of Botany, Zámek 1, Průhonice CZ-25243, Czech Republic; Department of Botany, Faculty of Science, Charles University, Benátská 2, Prague CZ-12801, Czech Republic; Department of Botany, University of Innsbruck, Sternwartestraße 15, 6020 Innsbruck, Austria; Prague Botanical Garden, Trojská 800/196, Prague CZ-17100, Czech Republic; Department of Experimental Plant Biology, Faculty of Science, Charles University, Viničná 5, Prague CZ-12844, Czech Republic

**Keywords:** orchids, diversification, partial endoreplication, evolution, Hyb-Seq, Pleurothallidinae

## Abstract

Some of the most burning questions in biology in recent years concern differential diversification along the tree of life and its causes. Among others, it could be triggered by the evolution of novel phenotypes accelerating diversification in lineages that bear them. In the Pleurothallidinae, the most species-rich subtribe of plants on Earth with 46 genera and ∼5,500 species, we constructed a completely new phylogeny and mapped on to it the type of endoreplication intending to trace how the phenomenon of partial endoreplication, which is unique to orchids, affects the differential diversification of lineages.

We have used NGS based target enrichment HybSeq approach for the reconstruction of the phylogeny and the flow cytometry to estimate the type of endoreplication. The BAMM and BiSSE analyses have been used to assess diversification rates and to trace the phenotype changes.

We have found that three of six changes in diversification rates are associated with changes in the endoreplication type and the clades bearing taxa with partial endoreplication showed higher net diversification rates. Our results demonstrate that multiple evolution of partial endoreplication within the subtribe considerably shapes the patterns of diversity and that partial endoreplication is a trait with an evolutionary significance.

## Introduction

Species richness is unequally distributed across the tree of life and remarkable differences are apparent even between closely related evolutionary lineages. In orchids, which are perhaps the most diverse plant family (rivalled only by the Asteraceae), there are several examples of disproportionally diversified species assemblages spanning from ancient monogeneric lineages (e.g. the tribes Codonorchideae, with two species, or the Thaieae, with a single species) persisting for dozens of millions of years undiversified [1,2] to clades with relatively recent massive diversification producing thousands of species (e.g. the largest plant subtribe Pleurothallidinae) [3]. The evolution of novel phenotypes triggering increased net diversification rates is often suggested as a driver of such disproportion, despite relatively rare evidence of such phenotypes. In orchids, accelerations of net diversification have been correlated with the evolution of pollinia, the epiphytic habit, CAM photosynthesis, tropical distribution and pollination by lepidopterans or euglossine bees [1]. Nevertheless, no such trait is confined to the orchids. By contrast, the phenomenon of partial endoreplication, so far known only from orchids, has never been tested for its diversification significance.

Endoreplication, the process of intraindividual polyploidization usually connected with cell differentiation, is unevenly distributed across the plant kingdom and is ubiquitously observed only in a few plant families such as the Brassicaceae or Cucurbitaceae (see Leitch & Dodsworth for an overview) [4]. Conventionally, endoreplication involves the doubling of the whole nuclear DNA content without cell division, resulting in the formation of endopolyploids. The exception from this rule are orchids, in which there exists so-called partial endoreplication (PE) [5,6], where a species-specific fraction of nuclear DNA is replicated, resulting in nuclei with (substantially) less than two-fold increase in DNA content [7]. It has to be noted that only one type of endoreplication has so far been observed in each species, so there is no evidence of any intra-specific switch from one type to another. Partial endoreplication is known from all subfamilies of orchids except one [8] and exhibits clade-specific behaviour with uneven distribution along the orchid phylogeny [7,9]. The biological significance of endopolyploidy remains elusive and, in plants, is usually attributed to the ability to grow in extreme or variable conditions [10,11]. Gene expression, enhanced by endoreplication and modulated under various environmental conditions, might be a factor that likely lies behind it [12]. Nevertheless, evidence that conventional and partial endoreplication differ in their biological significance is still lacking. All data gathered to date indicate a prevalence of PE in orchid subfamilies where it has once evolved [7], except for the species-richest subfamily Epidendroideae, which includes ∼ 80% of genera with exclusively the conventional type of endoreplication (CE; our survey). Within the Epidendroideae, only the monophyletic subtribe Pleurothallidinae combines exceptional diversification and almost equal representation of PE and CE.

Pleurothallidinae is a young group restricted to tropical America which has recently undergone an extraordinarily rapid diversification, resulting in more than 5,500 currently accepted species [13,14]. Species of this group colonized various habitats from lowlands up to nearly 4,000 m a.s.l. The Andean orogeny has been identified as an important factor in the diversification of the Pleurothallidinae [3], and high variability in pollinator adaptation has been hypothesized to also drive the speciation [15]. However, the specific factor generating this diversification, which is exceptional among other Andean plant groups and is asymmetric, remains elusive, and the question is whether propagation of partial endoreplication within the subtribe might be such a factor.

To investigate this, we first constructed a robust phylogeny of the Pleurothallidinae based on the Hyb-Seq approach utilizing orthologous low-copy nuclear (LCN) loci for phylogenetics in non-model organisms [16]. Next, we dated the resulting species tree and reconstructed ancestral states for both PE and CE to determine the shifts between them within the subtribe. Simultaneously, we used the same tree to detect shifts in net diversification rates. Finally, we compared the results to evaluate the association between diversification rates and the type of endoreplication. Under the assumption of such an association, we tested the hypothesis that a shift in the type of endoreplication precedes each shift in diversification rates and that clades with partial endoreplication exhibit greater diversification rates.

## Material & Methods

### Taxon sampling

We sought to sample as many morphological groups and previously recognized lineages as possible. Because living tissues are required for the estimation of the type of endoreplication, we were unable to use herbarium specimens and were limited by the availability of living plants in our collection. Finally, we sampled 40 out of 46 recognized genera that were represented by 341 specimens of 337 species, plus seven species of outgroups. Because the generic concept of Pleurothallidinae changes relatively frequently, we mostly follow the concept of Karremans [13]. *Kraenzlinella* and *Dondodia* are included in *Acianthera* following Karremans et al. [17]. *Trichosalpinx* is treated in its strict sense following Bogarín et al. [18]. Morphologically unusual species, which have sometimes been treated as monospecific genera but without molecular phylogenetic evidence (*Andreettaea, Madisonia*), are classified as synonymous with *Pleurothallis*, following WCSP [14]. The adopted concept of genera of the Pleurothallidinae supplemented with the number of recognized species [14] is provided in Table S1, the complete list of analysed species in Table S2.

### DNA extraction

Total genomic DNA was extracted from 0.5 g of silica-dried leaf material via the Sorbitol method [19] with two modifications: 1,600 μl of Extraction Buffer per sample were used (instead of 1,300 μl) and 6 μl of RNase instead of 4 μl in the first step. The quality of the DNA was checked on 1% agarose gel and using a Qubit 2.0 fluorometer (Invitrogen, Carlsbad, CA, USA).

### Probe design

To select loci that are nuclear, single-copy and have orthologs across the Pleurothallidinae for sequence capture (HybSeq), we utilize our genomic data for *Stelis pauciflora* (P022) and the transcriptome for *Masdevallia yuangensis* (ID JSAG) from the 1,000 Plants (1KP) initiative (http://www.onekp.com/samples/single.php?id=JSAG). For these two species from different genera of Pleurothallidinae, we expected to find loci that will have likely orthologs also throughout the entire subtribe. To select loci, we utilized the marker development pipeline Sondovac [16] as described in detail in Results S1. One probe set targeting a total of 4,956 nuclear DNA loci was prepared using MYbaits (MYcroarray Ann Arbor, MI, USA). Biotinylated RNA probes were 120 bp long with 2× tiling density over target sequences. Additional checks were performed to eliminate probes targeting multi-copy loci by probe manufacturer.

### HybSeq Library preparation

For each sample, an enriched genomic library was prepared following the protocol of manufacturer as described in detail in Results S1. Twenty-four samples per library were pooled in equimolar ratios (overall, 18 libraries were prepared). In-solution sequence capture of 4,956 exons from 1,200 nuclear low-copy genes designed as described above was done using MYbaits custom probes (MYcroarray, Ann Arbor, MI, USA) following the manufacturer’s protocol (see details in Results S1). Hyb-Seq libraries were sequenced on an Illumina MiSeq instrument at OMICS Genomika (DNA sequencing laboratory of the Biological Section of Charles University), Biocev, Vestec, using 300-cycle kit (v.3, Illumina, Inc., San Diego, CA, USA) to obtain 150 bp paired-end reads.

### Processing of raw reads and species tree reconstruction

We used the HybPhyloMaker v.1.6.4 pipeline [20] for raw read filtering, mapping of the filtered reads to a pseudoreference (exon sequences used for probe design, separated by strings of 400Ns) and construction of gene alignments (see details in Results S1). Also gene trees and the species tree were reconstructed according to the HybPhyloMaker pipeline. Gene alignments with more than 30% of missing data and less than 100% species presence were excluded from further analyses. Alignment characteristics were calculated using AMAS [21], trimAl 1.4 [22] and MstatX (github.com/gcollet/MstatX). Gene trees were reconstructed using RAxML 8.2.4 [23] with 1,000 rapid bootstrap replicates (BS) and the GTRGAMMA model. Summary statistics were generated using AMAS [21] and plotted using R 3.2.3 [24]. The multispecies coalescent model implemented in ASTRAL 5.6.1 [25] was used to construct species trees with default settings. For usage in subsequent analysis (PhyParts), an additional species tree was constructed with ASTRAL 5.6.1 from gene trees with < 50% BS collapsed branches into a polytomy created in TreeCollapserCL 4 [26]. Additionally, alignments of all low-copy genes were concatenated and analysed by taking the supermatrix approach using FastTree 2 [27]. An overview of the reads obtained for all taxa involved is given in Table S2.

### Gene tree species tree (in)congruence

We evaluated the level of concordance among gene trees (with a maximum of 30 percent of missing data and with collapsed nodes with < 50 % support) against the ASTRAL species tree with the program PhyParts (https://bitbucket.org/blackrim/phyparts) [28]. The species tree was previously rooted in Figtree [29] with representatives from subtribe Laeliinae as an outgroup. The output obtained with PhyParts was visualized by plotting pie charts on the ASTRAL species tree and ML concatenated tree with the script PhyPartsPieCharts (https://github.com/mossmatters/MJPythonNotebooks) using the ETE3 Python toolkit [30].

### Temporal scaling of the species tree

A few unambiguous orchid macrofossils are available for Orchidaceae, e.g. [31,32], but these are assigned to lineages very distantly related to the Pleurothallidinae. Because using distant outgroups to calibrate our phylogenies would have created extensive sampling heterogeneities, which can result in spurious results [33], we had to rely on secondary calibrations. As secondary calibration points we relied mainly on a previously fossil-calibrated Orchidaceae-wide phylogeny; in particular we used diversification time of the subtribe Pleurothallidinae to be ca 20 million years ago [3]. For the large genomic datasets similar to ours, the use of Bayesian methods for dating analyses (e.g. BEAST) [34] is frequently unfeasible, as they require a long computational times [35–38]. Here we used the RelTime method [39], a fast-dating and high-performance algorithm that is implemented in MEGA X [40]. We used a species tree rooted with representatives of the subtribe Laellinae as a starting tree and concatenated alignment of full sequences of all 168 nuclear loci as source data for branch length estimation. The GUI-based version of RelTime in MEGA X was utilized, using the RelTime-ML option for timetree estimation and GTR with four discrete gamma rate categories as a model for the sequence alignment. As a time constraint, we used prior information on the diversification of the Pleurothallidinae, specifically 20 ± 5 Mya [3].

### Endoreplication mode estimation

Data on endoreplication mode (partial vs conventional endoreplication) were estimated via flow cytometric analyses of all samples included in the tree according to the methodology described in detail by Trávníček et al. [8]. This approach is based on the quantification of the amount of DNA that has undergone endoreplication in the course of cell differentiation. Whereas plants with 100% replicated DNA (endoreplicated nuclei of subsequent sizes always differ two-fold) are assigned to the conventional type of endoreplication, plants with less than 100% replicated DNA (endoreplicated nuclei of subsequent sizes always differ by less than two-fold) are assigned to partial endoreplication (Fig. S1). The arbitrary threshold for assignment was set to 95% of replicated DNA because of possible measurement errors. The assignment of all included taxa was conducted via FloMax v2.4d (Partec GmbH) and is given in Table S2. In addition to the assignment of all involved ingroup plants (341 individuals), 340 additional plants randomly distributed along an ingroup clade were assigned to on of the endoreplication types to increase the precision of the estimation of the proportion between the two types of endoreplication within the Pleurothallidinae.

### Ancestral state reconstruction

Using the species tree, we reconstructed ancestral states for endoreplication type under a BiSSE model (binary state speciation and extinction model) [41] implemented in the R package *diversitree* [42]. Incomplete sampling was accounted for by a state-dependent sampling factor that was estimated by the use of an extended dataset (see above). We started with a full six-parameter model and performed model simplification in a maximum likelihood framework based on likelihood ratio tests (implemented in the R package *lmtest*) [43]. Two best-fitting five-parameter BiSSE models with equal likelihood were found: the first model with single extinction rate and the second with a single speciation rate. Initially, the reconstruction using stochastic character mapping was performed using the R package *phytools* v.06-99 [44] and the function make.simmap. This method fits a continuous-time reversible Markov model for the evolution of the trait and then simulates stochastic character histories using that model and the tip states on the tree (sensu Bollback) [45]. We simulated 1,000 character histories for the endoreplication trait and summarized it (appropriate nodes in Fig. **1**; all nodes in Fig. S2). We also performed ancestral-state reconstruction for the BiSSE models (Figs. S3–5).

**Figure 1.**
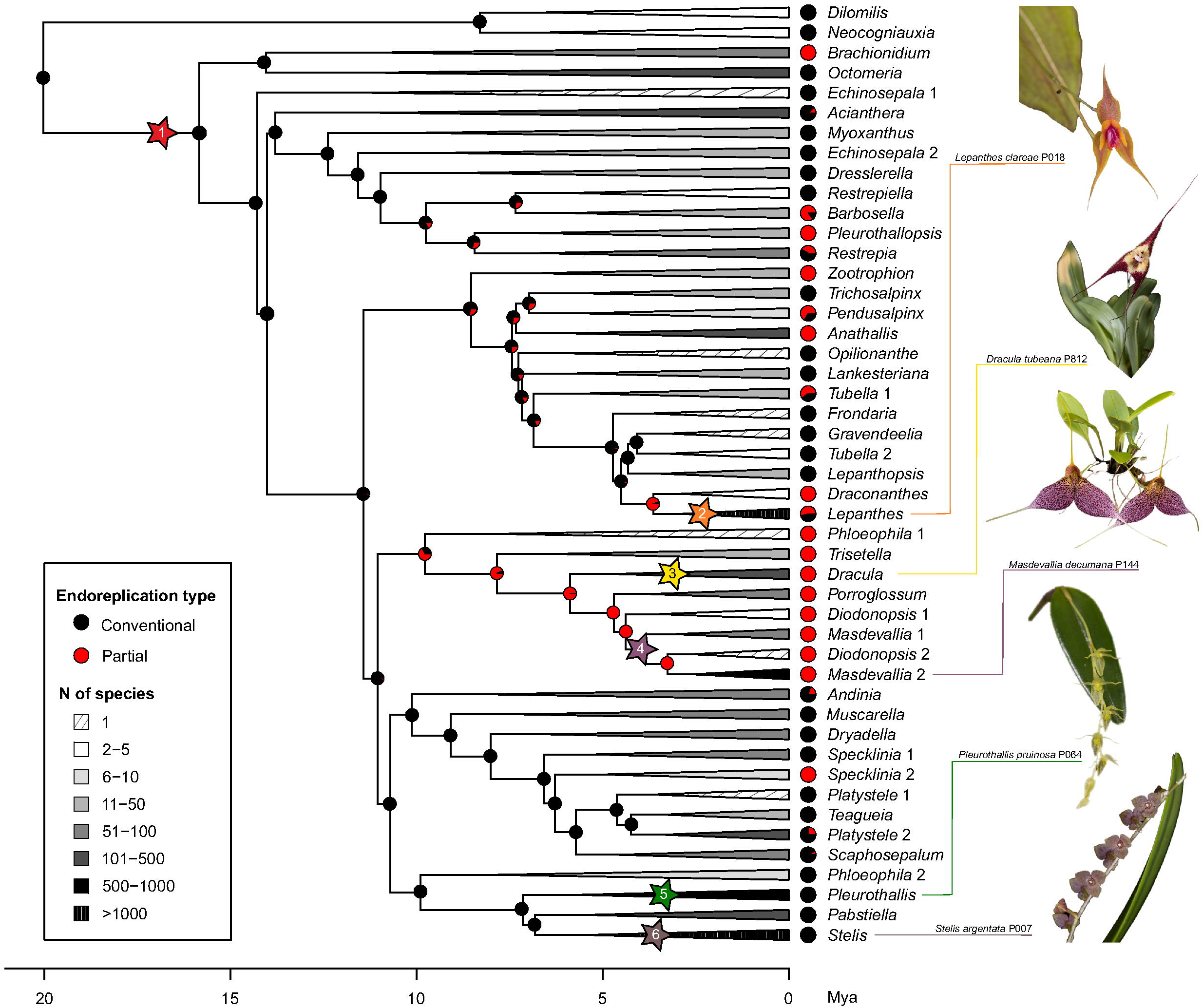
Diversity tree for analyses of lineage diversification within the Pleurothallidinae based on a simplified species tree obtained using the ASTRAL approach. Clades are collapsed to 47 monophyletic lineages representing 40 recognized genera and coloured by extant species diversity. Clades with significantly increased net diversification rates are labelled by numbered and coloured asterisks. Clades are supplemented with pie charts reflecting the proportion of species based on their endoreplication type. Nodes are provided with pie charts reflecting an ancestral state reconstruction derived from stochastic character mapping of endoreplication type on the species tree.

### Diversification rates

We used Bayesian analysis of macroevolutionary mixtures, BAMM v2.5.0 [46] to estimate the pattern of diversification within the Pleurothallidinae. The model allows for variation in rates over time and among lineages, and despite criticism that it uses an incorrect likelihood function and prior settings [47], it provides a robust and appropriate tool for estimating lineage diversification [48,49]. BAMM allows for the implementation of incomplete clade-specific taxonomic sampling, which was necessary because of the use of a phylogeny reflecting just a fraction of the species diversity of the Pleurothallidinae (341 individuals in the phylogeny vs ∼ 5,500 recognized species). We therefore applied fractioning of recognized diversity according to clade membership and virtually attribute each tip within a clade to represent the proportional fraction of known clade species richness (Table S1). We performed multiple BAMM runs of 50 millions generations, sampling every 2,000 generations, and checked the convergence using the R package *coda* v0.19-2 [50]. The effective sample size (ESS) of the log-likelihood and the number of shift events present in each sample remarkably exceeded the recommended threshold of 200. The analysis was repeated three times to ensure the convergence of all runs. BAMM output data were further processed using the R package *BAMMtools* v2.1.7 [51] that enables complex analysis and graphical representation.

### Diversification vs trait correlates

We estimated speciation, extinction and net diversification rates for clades according to the type of endoreplication – PE, CE and mixed clades. We extracted posterior distribution of state-dependent rates from BAMM analyses and examined the differences in rates between clades. For this we used the getCladeRates function of *BAMMtools* to obtain mean rates for particular clades. We compared state-dependent rates via credible intervals of differences to assess their significance. Similarly, we examined the best BiSSE model (with equal extinction rates) using a MCMC chain of 10,000 generations. We set the exponential prior to 1/2*r*, where *r* was the character-independent diversification rate inferred from the BiSSE starting point. We removed 10% a burn-in and summarized MCMC samples to assess the variation in state-dependent rates. Finally, we calculated differences in posterior distributions of rates for character-states (PE vs CE) and estimated their credible intervals for the best (i.e. that having equal extinction rates) and the full BiSSE model (Fig. S6). In the last step, we compared the results derived from the BAMM and BiSSE approach.

### Relative flower size

We recorded simple morphological traits (length of the ramicaul, length and width of leaves, and length of sepals and petals) from the appropriate literature sources to estimate the relative size of generative organs (flowers) to that of vegetative part of plants. We excluded taxa with unclear species determination. Those traits were investigated with respect to the type of endoreplication to elucidate the possible impact of PE on the morphological configuration of plants under investigation. We gathered data for 322 taxa (Table S2) and analysed them by ANOVA with and without phylogenetic independent contrasts (PIC) in the R package *phytools* [44] and *stats*, respectively.

## Results

### Phylogenetic and dating analyses

Species tree topologies obtained via the ASTRAL method and the supermatrix approach (concatenation) based on 168 loci differed only slightly, both showing high support values of Bayesian posterior probabilities (PP) in the case of the ASTRAL tree and SH-aLRT (SH) in the case of the supermatrix approach (Fig. S7). The large size and complexity of genomic datasets for phylogenetic reconstruction introduce significant phylogenetic noise and conflict into subsequent analyses. For such datasets the Bayesian posterior distributions, SH-like local supports and even bootstrap supports are less informative because usually only a small minority of nodes show lower than maximum support, and the values might be biased by the size of the supermatrix [28,52]. Therefore we also examined the number of gene trees that are in concordance and in conflict with the species tree topology to interpret the support for particular phylogenetic units as well as to identify potential sources of conflicts like hybridization or incomplete lineage sorting [28]. PhyParts analysis of 168 gene trees and the ASTRAL species tree (Fig. S7a) shows high support (more than 50% of gene trees, hereafter GT, that support the species tree topology) for monophyletic Pleurothallidinae (160 GT) as well as for the core of the subtribe after separation of the basal genera *Dilomilis* and *Neocogniauxia* (144 GT). Support for particular genera and groups of genera, however, differs markedly across the phylogeny as described thoroughly in Methods S1. In addition, the groups of our interest, that is, the genera where shifts in diversification rates were observed (see below), differed in the number of gene trees that supported their monophyly, although the posterior probability was in all cases 1. In particular, the genus *Lepanthes* had the highest proportion of concordant/conflicting gene trees – 147/1. In the genera *Pleurothallis, Dracula* (excluding the potentially hybridogenous species *D. xenos*, see details in Methods S1) and *Stelis*, proportions of concordant genes were lower but still greater than those of conflicting ones (90/12, 60/29 and 40/32, respectively) and for the genus *Masdevallia* (including one species of the polyphyletic genus *Diodonopsis*) the proportion of conflicting gene trees exceeds that of concordant ones (28/47). The first clade that includes the genus *Masdevallia* and has a greater proportion of concordant genes (65/29) also covers the genera *Porroglossum* and *Diodonopsis*.

The absolute ages obtained for Pleurothallidinae chronograms are in agreement with previously published dated phylogenies, e.g. [3,53–55]. Divergence time estimates and 95% credibility intervals (CIs) inferred for all nodes of Pleurothallidinae chronograms are presented in Fig. S8. For the sake of clarity, we present the results using a simplified dated species tree visualizing major clades that roughly correspond to the accepted concept of genera. We divided clades with the incidence of paraphyletic genera into monophyletic lineages and supplemented the genus names with numbers to distinguish them (Fig. **1**). Whereas the divergence time of the ingroup (Pleurothallidinae) was set as prior to 20 mya (± 5 mya), the age of the subtribe after the separation of the basal genera *Dilomilis* and *Neocogniauxia* (i.e. diversification of the Pleurothallidinae) was estimated to be 15.8 (CI 9, 24) mya. Below we present divergence times of groups of our interest along with information about their diversification rates.

### Modelling diversification rates

To obtain diversification patterns within the phylogenetic tree, we used Bayesian analysis of macroevolutionary mixtures (BAMM) [46]. The analysis revealed multiple shifts in diversification rates in the Pleurothallidinae (Fig. **1**, Fig. S9). We found strong support for 4– 11 remarkable shifts in diversification rates across the subtribe, with a cumulative posterior probability of 0.98. The best models generated by Bayesien credible sets of shifts consistently show the six most preferred shifts (Fig. **1**, Fig. S10). The first shift is located at the basis of the subtribe after the separation of the basal genera *Dilomilis* and *Neocogniauxia* (ca 15.8 mya), the other five shifts are more or less confined to the genera *Lepanthes, Dracula, Masdevallia, Pleurothallis* and *Stelis*, respectively. Whereas the model identified *Lepanthes* and *Masdevallia* as participating in increased diversification as whole genera, other genera are involved partially. The genus *Dracula* without the controversially placed *D. xenos*, the genus *Pleurothallis* without numerous basal lineages and genus *Stelis* only in its nominate subgenus *Stelis* (Fig. S9). Mean divergence times of the clades with increased diversification rates are generally not older than 4 million years (see details in Table S3), where group of *Masdevallia* 1 & 2 (incl. *Diodonopsis* 2) seems to be the oldest one (mean 3.8 mya, 95% CI 1–7.6 mya). In accordance with common practice [56], we also computed the branch-specific marginal probability and marginal odds (branch-specific pseudo-Bayes factors) to visualize the branches with rate shift locations. Both approaches show consistent results that are in accordance with the location of preferred shifts (Fig. S11). We also performed an analysis of diversification rates over time to reveal clade-specific rates in contrast to rates of the background (the rest of the Pleurothallidinae excluding the particular clade; Fig. **2**). Those data show contrasting patterns in net diversification rates, diverging by the type of endoreplication. Whereas rates for clades with partial endoreplication (Fig. **2c** and **2d**) increase constantly, in accordance with the background, rates for clades with conventional endoreplication (Fig. **2e** and **2f**) exhibit a decrease. The exceptionally high net diversification rate for *Lepanthes* (Fig. **2b**), the only diversified clade with mixed endoreplication, is constant over time.

**Figure 2.**
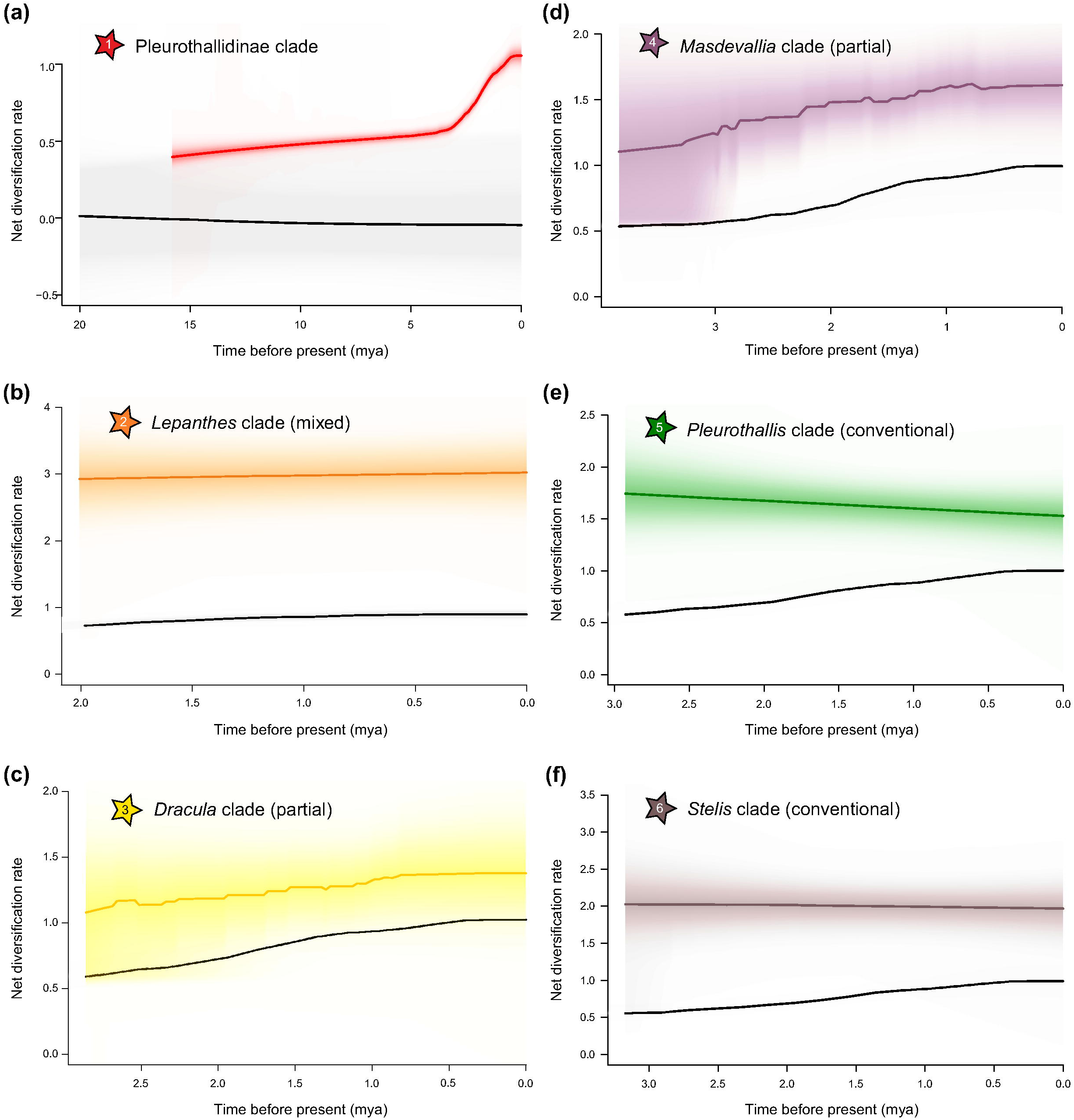
Net diversification rates over time for the six clades with consistent and distinct shift configuration inferred from BAMM analysis. The graphs correspond to the clades marked in Fig. **1** and represent different types of endoreplication: (a) a clade of almost all Pleurothallidinae except two basalmost lineages, (b) a clade of the genus Lepanthes with the mixed type, (c) a clade of the genus *Dracula* with the partial type, (d) the genus *Masdevallia*, (e) a clade of the genus *Pleurothallis* with the conventional type, and (f) the genus *Stelis*. Coloured lines indicate net diversification rates and black lines represent net diversification rates of the background phylogeny. Shading around lines is for 90% confidence intervals.

### Ancestral state reconstruction and diversification

We used ancestral state reconstruction via BiSSE and stochastic mapping to trace the possible shifts in the type of endoreplication and their possible coincidences with shifts in diversification rates. Assuming the significance of endoreplication in the diversification of orchids, we hypothesized that a shift in the type of endoreplication type preceded a shift in diversification rates. Such a pattern is traceable namely in the clade *Draconanthes*/*Lepanthes*, where massive diversification (Fig. **2b**) immediately follows the shift to partial endoreplication or to a mixture of both types (Fig. S2). Other shifts of diversification rates in the *Dracula* and *Masdevallia* clades follow the shift to exclusively partial endoreplication with a noticeable delay (Fig. S2). The remaining increases of net diversification in *Pleurothallis* and *Stelis* do not show any association with shifts in endoreplication type because they are confined to clades with exclusively CE (Fig. **1**, Fig. S2).

### Trait-dependent diversification rates

Finally, we estimated trait-dependent diversification rates in clades with differential presence of endoreplication types. We primarily used the approach of taking posterior distribution of state-dependent rates from BAMM analyses. The clades were divided into three groups according to endoreplication type: CE clades containing only taxa with the conventional type, PE clades with exclusively the partial type and mixed clades where taxa exhibit either the conventional or the partial type of endoreplication. The clades were treated at the level of genera, but reflecting monophyletic lineages with the exception of the genus *Acianthera*, which was a priori divided down to the basalmost lineage corresponding to the subgenus *Kraenzlinella* (PE clade) and the rest of species (CE clade). Under this treatment we revealed significant shifts towards increased net diversification and speciation rates in PE and mixed clades; however, extinction rates remained almost the same for all clades (Fig. **3**). Significance was assessed by examining the posterior distribution of differences between the clades (Fig. S12). Alternatively, we used posterior probabilities from the fit of the BiSSE model, where only two states of the trait were recognized (PE vs CE) because the analysis was carried out at the species level. The BiSSE model with equal extinction rates shows an even more prominent shift of net diversification rates for taxa with PE (Fig. S13). Additionally, we tried two alternative setups of clade partitioning for the interpretation of state-dependent rates from BAMM. We analysed the data at the level of whole clades (without dividing the genus *Acianthera*) and, conversely, at the finest (species) scale, where the assignment to mixed clades was limited only to sister taxa with a different type of endoreplication. Whereas the first approach changed noticeably, the estimation of net diversification rates for mixed clades, the second approach provided similar results as previous analyses based on BAMM and BiSSE data (Fig. S14).

**Figure 3.**
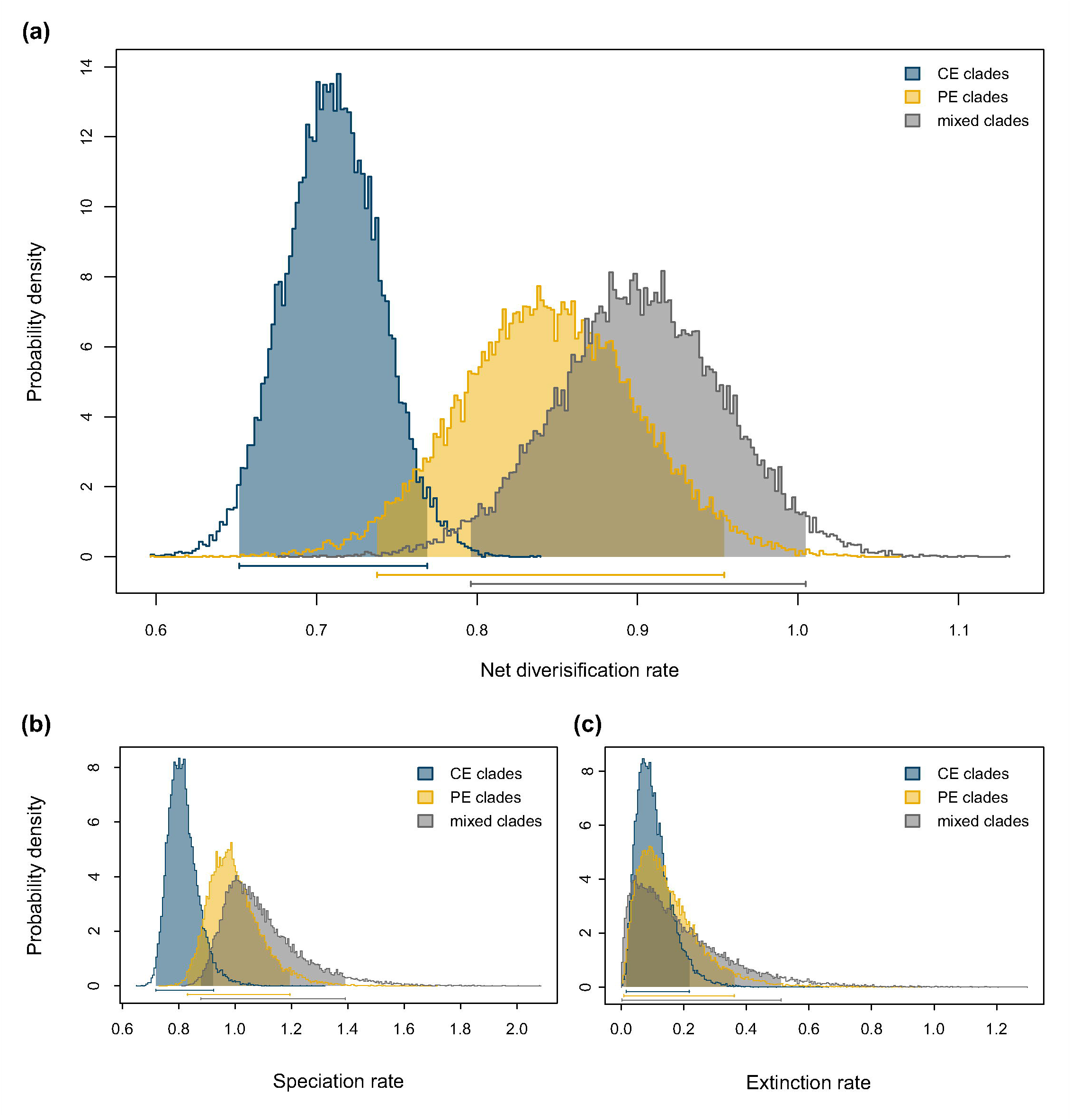
State-dependent diversification rates extracted from BAMM analyses corresponding to three types of clades according to the type of endoreplication present – clades with exclusively conventional endoreplication (CE clades), clades with exclusively partial endoreplication (PE clades) and clades with both types (mixed clades). The graphs show the posterior distribution of rates for (a) net diversification, (b) speciation and (c) extinction, coloured by character state.

### Morphological traits

We also tested the possible impact of PE on the phenotype to trace the importance of saving energy in connection with morphology. For this, we made a simple comparison of vegetative and generative parts of the Pleurothallidinae within our phylogeny and analysed data with respect to the type of endoreplication (Table 1). Flower size (represented by the length of the sepals), in both an absolute and relative way, was singled out as the most prominent difference between taxa differing in the type of endoreplication even when the data were tested under the phylogenetic frame.

**Table 1.** Analysis of variance (ANOVA) for selected morphological traits adopted from the literature with respect to the type of endoreplication (pairwise comparison of PE and CE taxa). P values both for ordinary and phylogenetic ANOVA are provided and values in bold are statistically significant (α - 0.5). Arrows next to p-values indicate the direction of trait shifts for PE taxa.

| Trait | F value | p<sub>ANOVA</sub> | p<sub>phyloANOVA</sub> |
| --- | --- | --- | --- |
| Ramicaul length | 18.255 | < **0.001** ↓ | 0.364 |
| Leaf length | 2.724 | **0.010** ↑ | 0.717 |
| Leaf width | 0.079 | 0.779 | 0.954 |
| Leaf size (length x width) | 0.186 | 0.667 | 0.937 |
| Sepal length | 87.339 | < **0.001** ↑ | **0.038** |
| Petal length | 0.7 | 0.404 | 0.863 |
| Sepal length / leaf size | 13.5 | < **0.001** ↑ | 0.457 |
| Sepal length / ramicaul length | 62.366 | < **0.001** ↑ | 0.08 |
| Sepal length / ramicaul + leaf length | 84.558 | < **0.001** ↑ | **0.037** |

## Discussion

We present here the first thorough phylogenetic study of the whole Pleurothallidinae based on NGS data, using own designed probes for target enrichment according to [16]. Our approach revealed relatively high support for the current taxonomic concept in many lineages (genera), but indicated also paraphyly in some salient examples such as *Masdevallia*-*Diodonopsis, Tubella, Phloeophila, Specklinia, Platystele* and *Echinosepala*. The latter findings are either new or contradict the latest phylogeny of the subtribe inferred by [3] based on ITS and *mat*K. We also present a relatively critical approach to the interpretation of phylogenies based on genomic data while taking into account numbers of gene trees that are concordant and discordant with the species tree reconstruction. We see multiple indications of hybridogenous origins of species or entire lineages (e.g. species of the genera *Myoxanthus, Restrepia, Zootrophion, Tubella, Lepanthopsis, Dracula, Specklinia, Stelis* and several others). Reticulate evolution has been documented also in several other orchid groups like *Paphiopedilum* (subfamily Cypripedioideae) [57], *Dactylorhiza* (subfamily Orchidoideae) [58] and *Polystachya* (subfamily Epidendroideae) [59], which may indicate that this process plays an important role in orchid evolution. In parallel we see indications of rapid radiations accompanied by massive gene tree – species tree incongruence likely caused by incomplete lineage sorting (e.g. in the genera *Masdevallia* and *Stelis*). Similarly high proportions of conflicting gene trees have been observed in young lineages of the rapidly diversified genus *Picris* [60]. We believe that our approach to thoroughly evaluating patterns of gene trees that conflict or support the topology of a species tree will be valuable for further detailed systematic/taxonomic studies, where the above mentioned evolutionary phenomena need to be taken into account, as shown, for example, by [61].

Possible weaknesses of our research stem from the nature of the subtribe under study and the impossibility to account for all recognized species, of which there are more than 5,500 [13,14]. Despite low coverage of the subtribe in our phylogeny, accounting for ∼6.2%, we are convinced of the reliability of our results because we covered almost all out of the 46 recognized genera (except for six very small genera; Table S1). A very similar approach including a near-complete generic level encompassing only a fraction of species was taken by Helmstetter et al. [62]. Our conviction is supported by convergent results from two completely independent methods with different rationales and handling with incomplete sampling. In addition, our phylogeny fulfils basic recommendations for BiSSE-like analyses to avoid the main pitfalls of low sample size and strong tip ratio bias [63,64].

The time coincidence of massive acceleration in the five most diverse lineages (∼2.4–3.8 Ma) points to the existence of a common, probably an environmental, trigger. The establishment of suitable habitats after the main orogenesis of the Andes has been postulated as the most reliable explanation [3]. However, clade-specific net diversification rates exhibit substantial differences (Fig. **2**), suggesting the existence of other factors differentiating the clades. Under this assumption, we tested the possible impact of partial endoreplication on divergence within clades. We have found three of five shifts in diversification rates related to genera associated with a shift from conventional to partial endoreplication. In addition, our data unequivocally imply an association between greater diversification rates and PE clades (Figs. **2** and **3**). So, how might partial endoreplication promote diversification in orchids? PE might be enhanced by the necessity to have an increased gene expression, crucial for plants with fast expansion growth of cells. Such growth is not attained by increasing the ploidy level of nuclei, but by regulating the expression of cell wall-modifying genes that allow cell growth via vacuolar expansion [65]. Therefore, endoreplication might be of particular importance for tissues containing extremely rapidly expanding cells, where an increase in the gene copy number through endoreplication might be a way to cope with the high demand for new cell wall material. The possible advantage of PE over CE might lie in its relatively low energetic and supply costs because increased expression can be facilitated by endoreplication of just a fraction of the genome [9]. This might be important especially in extreme environmental conditions or under strong competitive pressure, that is, in places the majority of the Pleurothallidinae grow. We can further speculate that the energy saved by partial endoreplication might be invested into the formation of more attractive flowers for pollinators, so PE might serve as a trigger of increased diversification via speciation driven by pollinator adaptation [15]. According to this assumption, genera with the largest flowers within the subtribe (*Dracula* and *Masdevallia*) perform only PE whereas genera known for relatively large plants with small flowers (*Myoxanthus, Octomeria, Pleurothallis, Stelis*) perform only CE. Moreover, several other genera with a high incidence of PE are tiny plants, where the size of the flower is relatively large compared to that of vegetative parts of the plant (*Anathallis, Brachionidium, Lepanthes*). We therefore decided to analyse relative proportions of flowers to vegetative parts in sampled species of the Pleurothallidinae, and the results of this analysis clearly show that species with PE can invest more resources into producing larger flowers (Table 1). Even though the role of large flower size in myophilous (i.e. fly-pollinated) species, a substantial fraction of the Pleurothallidinae, remains a mystery, it has been observed in the genera *Restrepia* and *Telipogon* that osmophores can be distributed on various parts of flowers, sometimes very far from each other, likely to produce a more intensive gradient of a scent or to make ‘scent trails’ to guide pollinators [66,67]. We could therefore hypothesize that large flowers might provide an evolutionary advantage also in myophilous species of the Pleurothallidinae. However, there is another completely different, yet also plausible, explanation for the dominant diversification of PE clades. Endoreplication is induced in mycorrhizal tissues by fungal accommodation in arbuscular mycorrhizas [68] as well as in orchid mycorrhizas [69,70]. Taking into account that orchids are completely dependent on nutrition mediated by mycorrhizal fungi at the (post-) germination stage [71,72], we can deduce that they are predetermined to deal with endoreplication. We can therefore speculate that partial endoreplication might be a way to tackle the predetermination fostering its advantages while minimizing its disadvantages by saving resources also during mycorrhiza-induced endoreplication. Taking into account that PE is prominent in orchid species with larger genome sizes [7], we can further speculate that PE might represent an effective evolutionary response on how to simultaneously cope with limited sources and endopolyploidy. Seen in this light, the greater diversification rate in clades harbouring PE makes perfect sense. Anyway, elevated diversification rates are not exclusively associated with shifts in endoreplication type within the Pleurothallidinae, but clades with partial endoreplication exhibit greater rates in direct comparison to clades with conventional endoreplication (Fig. **3**). To sum up, we can conclude that PE plays a noticeable role in the evolution of the Pleurothallidinae and contributes to the formation of the species-richest subtribe of plants on Earth.

## Conclusions

Our results demonstrate that the largest orchid subtribe Pleurothallidinae diversified extremely rapidly in the last ∼ 4 Mya through frequent hybridogenous speciation accompanied by the multiple evolution of partial endoreplication, a feature unique to orchids. The five most species-rich lineages radiated at an increased rate around the same time, which points to the existence of a common, probably environmental, trigger. In addition, the evolution of partial endoreplication strongly correlates with the diversification of clades possessing it. The full evolutionary consequences of partial endoreplication are yet to be understood, but species with PE save considerable resources by not performing full endoreplication and can invest them into producing larger flowers to better attract pollinators. Indeed, PE is more common in species with bigger flowers and larger genome sizes, so it might represent an effective evolutionary workaround reducing the cost of endoreplication in polyploids.

## Supporting information

Cover letter

## Acknowledgements

We are indebted to Blanka Hamplová and Štěpánka Hrdá from the BIOCEV centre for their help with sequencing and to F. Rooks for kindly improving the English.

## Author contributions

Z.C., E. Z., J.P. and P.-A.S. contributed equally to the work supervised by P.T. All authors carried out the analyses, interpreted and discussed the data and contributed to the writing and editing of the manuscript.

## Data availability statement

Sequence data that support the findings of this study have been available in a public repository deposited in GenBank with the primary accession code PRJNA604559. The flow cytometric data that support the findings of this study are available from the corresponding author upon reasonable request.

## Supporting information

Electronic supplementary material is available online.

**Results S1.** Extended Results

**Methods S1.** Extended Materials and Methods

**Fig. S1.** Flow-cytometric histograms and scatter plots for two species of the genus *Lepanthes* showing a different type of endoreplication.

**Fig. S2.** Ancestral state reconstructions of endoreplication type using stochastic character mapping.

**Fig. S3.** Ancestral state reconstructions of endoreplication type using a full BiSSE model.

**Fig. S4.** Ancestral state reconstructions of endoreplication type using a BiSSE model with equal extinction rates.

**Fig. S5.** Ancestral state reconstructions of endoreplication type using a BiSSE model with equal speciation rates.

**Fig. S6.** Credible intervals of differences from the posterior distribution of samples taken from BiSSE analyses.

**Fig. S7.** Species tree topologies obtained via the ASTRAL approach (a) and ML concatenation (b) of Pleurothallidinae.

**Fig. S8.** Divergence time estimates and 95% credibility intervals (CIs) inferred for all nodes of Pleurothallidinae chronograms.

**Fig. S9.** Mean phylorate plot of net diversification from BAMM analysis.

**Fig. S10.** The nine most common credible shift sets from BAMM analysis.

**Fig. S11.** Relative evidence of shifts in endoreplication mode inferred from the posterior distribution of BAMM results.

**Fig. S12.** Credible intervals of differences from the posterior distribution of samples taken from BAMM analyses and divided according to the type of endoreplication into partial (PE), conventional (CE) and mixed clades.

**Fig. S13.** Posterior distribution of state-dependent rates extracted from BiSSE analyses with equal extinction rates.

**Fig. S14.** State-dependent diversification rates extracted from BAMM analyses corresponding to three types of clades according to the harboured type or types of endoreplication – clades with exclusively conventional endoreplication (CE clades), clades with exclusively partial endoreplication (PE clades) and clades with both types (mixed clades).

**Table S1.** List of specimens included in the study supplemented by the type of endoreplication, morphological traits and HybSeq statistics (as separate Excel table).

**Table S2.** List of genera according to the adopted taxonomic concept supplemented with the number of species included, total species richness, endoreplication type, and data inferred from ASTRAL topology.

**Table S3.** Summary characteristics for clades corresponding to genera with significantly increased diversification rates.

