## Supplementary material for "Partial endoreplication stimulates diversification in the species-richest lineage of orchids": Cover letter

INSTITUTE OF BOTANY CAS

Průhonice, Czech Republic; May 12, 2020

Dear Editor,

please find attached our manuscript entitled '**Partial endoreplication stimulates diversification in the species-richest lineage of orchids**'. This is the first submission of the manuscript. We believe our study addressing the evolutionary significance of partial endoreplication in the diversification of orchids fits well within the scope of the journal, as is further explained below.

Cell differentiation in orchids is accompanied by endoreplication of two types, conventional (CE), where whole nuclear DNA is replicated, and partial (PE), where only a species-specific part of nuclear DNA is replicated. We have investigated whether PE, a pathway of cell differentiation unique to orchids, could act as a trigger of their extraordinary diversification. We have addressed this topic on the example of the species-richest monophyletic orchid subtribe Pleurothallidinae by a combination of BAMM and BiSSE modeling. For this, we constructed a completely new phylogeny of the subtribe using the HybSeq approach with a suite of targeted probes. Subsequently, we mapped the type of endoreplication on to the phylogeny to trace whether the transition from CE to PE is associated with the diversification of particular lineages.

We have found three out of five shifts in diversification rates to be related to genera associated with shifts from CE to PE. Moreover, our data unequivocally imply an association between greater diversification rates with clades where PE species outbalanced CE species.

Finally, we broadly discuss the possible avenues of how PE acts as an evolutionary advantage in orchids. Although the answer to this question remains elusive, we can conclude that PE plays a noticeable role in the evolution of the Pleurothallidinae and contributes to the formation of the species-richest subtribe of plants on Earth. This is the first time that partial endoreplication, a physiological trait operating at the cell level, which is of an unknown origin and has a not fully understood function, has been identified as a significant promoter of orchid speciation.

Thank you for considering this manuscript for publication.

Yours faithfully,

Pavel Trávníček, corresponding author
