## Supporting information for "Partial endoreplication stimulates diversification in the species-richest lineage of orchids"

**Methods S1.** Extended Materials and Methods

*Phylogenetic reconstructions based on a low-copy gene dataset.* We grouped the genera of *Pleurothallidinae* into six sections based on the topology of the ASTRAL tree and the concatenated species tree (Fig. S1a and S1b), and their support in either local posterior probability, SH-aLRT values or in gene tree/species tree concordance (based on PhyParts analysis). Here we consider clades well supported if the portion of concording gene trees is greater than the amount of conflicting gene trees. Into basal grade "0" we assigned 12 genera occurring in the basal position of the *Pleurothallidinae* whose mutual relationships as well as their relationships to the other ingroup genera are only poorly supported. The monophyly of all genera is, however, well supported with the exception of *Echinosepala*, which appears polyphyletic. Group "A" includes 12 genera, 106 gene trees supporting the species tree and 100% support in ASTRAL analysis. Ten genera are well supported whereas *Lankesteriana*, *Opilionanthe* and *Tubella*2 are weakly supported both in gene/species tree concordance and in the ASTRAL analysis. *Gravendeelia*, which itself is well supported, clustered together with weakly supported *Tubella*2, and this conflict on the higher order is reflected in different topologies between the ASTRAL and concatenated tree. Whereas *Lepanthes katleri* (P015) and *Lepanthes camprimulgus* (P271) are well supported, the internal support for *Lepanthes* is generally weak. *Trichosalpinx*, *Pendusalpinx* and *Anathallis*, which themselves are well supported, are clustering together and are weakly supported on the higher level. This is also reflected in the discordance between the ASTRAL and the concatenated tree. Group "B" is composed of five genera and supported by 141 gene trees and 100% support in the ASTRAL analysis. Well supported genera are *Poroglossum* and *Tristella*, weakly supported are *Diodonospsis*1 and 2 which appear polyphyletic while *Masdevallia* appears paraphyletic with *Masdevallia*1 being well supported in contrast to *Masdevallia*2. Though, the concatenated tree shows almost the identical pattern as the ASTRAL tree. *Dracula* is supported evenly by 51 gene trees concordant and conflicting with the species tree, respectively. The topologies of the ASTRAL and the concatenated trees show some minor differences here, highlighted by tangelgrams in Fig. S1 Group "C" only consists of *Andinia*, which is well supported with 145 gene trees matching the species tree, and 100% support in ASTRAL. Except *Andinia shizopogon* the support within *Andinia* is high. Group "D" includes six genera that are supported by 64 gene trees (32 conflicting) and 100% in ASTRAL. Weak support is seen only in *Specklinia*1. Yet, the topologies between the two analyses are concordant. The relations among *Scaphosepalum* and *Platystele*/*Teagueia* are weakly supported, as well as parts of internal relations in *Scaphosepalum* and *Platystele*2. The Group "E" is composed of three genera – *Pabstiella*, *Pleurothallis* and *Stelis*. This group has 111 gene trees supporting the species tree. The ASTRAL species tree shows 100% support in PP for this clade. *Papstiella* is well supported in both gene/species tree concordance and in the ASTRAL analysis. Yet, both *Pleurothallis* and *Stelis* are internally lacking support, in both ASTRAL analysis and gene/species tree concordance. This lack of support is also reflected by differences in topology between the two species trees. The genus *Phloeophila* is not assigned to a section in our results as it appears polyphyletic, the species *Phloeophila pelecaniceps* being located very basal compared to the species *Phloeophila pleurothallopsis* (samples P721 and P103).

**Results S1.** Extended Results

*Probe design*. To select nuclear, single-copy loci that have orthologs across the Pleurothallidinae, we utilized the marker development pipeline Sondovac [1]. Briefly, the pipeline takes genome read data (here we used genome skimming data from *Stelis pauciflora* P022) and removes any reads matching a plastome and reference. We used a plastome from *Masdevallia coccinea* (GenBank sample no. KP205432.1). Sondovac then removes duplicate transcripts from the transcriptome (here we used a transcriptome for *Masdevallia yuangensis* (ID JSAG) from the 1,000 Plants (1KP) initiative (http://www.onekp.com/samples/single.php?id= JSAG)) to identify single-copy loci and find genome reads matching the remaining unique transcripts, which are de novo assembled. Assembled contigs from genome reads are filtered for length (contig >120 bp, total length of all contigs for a transcript >600 bp) and uniqueness. Remaining contigs are compiled as target sequences. Sequences with >90% sequence similarity were identified using cd-hit-est [2], and the longest sequence in each cluster was retained.

One probe set (myBaits custom probes) targeting a total of 4,956 nuclear DNA loci was designed and manufactured by Arbor Biosciences (formerly MYcroarray, Ann Arbor, MI, USA). Biotinylated RNA probes were 120 bp with 2× tiling density over target sequences. Additional checks were performed to eliminate probes targeting multi-copy loci by the manufacturer.

*HybSeq – Library preparation*. Genomic DNA was fragmented using an M220 Focused-ultrasonicator (Covaris, Woburn, MA, USA; settings 52 s, 6 °C, 200 cycles), ~800 bp fragment length was verified using agarose electrophoresis. Library preparation followed the NEBNext Ultra DNA Library Prep Kit for the Illumina protocol (New England Biolabs, Ipswich, MA, USA), but the amount of all solutions was reduced by half, and purification was done using the QIAquick PCR Purification kit (Qiagen, Venlo, Netherlands). Agarose gel size selection (~500–600 bp) forwent the second purification step and followed the QIAquick Gel Extraction Kit (Qiagen, Venlo, Netherlands) protocol but with extended 15–30 min incubation at room temperature and 5 min elution to 10 µl $ddH_2O$. Libraries were enriched through 8 cycles of PCR with NEBNext Multiplex Oligos for Illumina Index Primers Set 1 (New England Biolabs, Ipswich, MA, USA) and twice cleaned up with Agencourt AMPure XP beads (Beckman Coulter Genomics, Danvers, MA, USA, ratios 0.7:1 and 0.65:1 beads : DNA) and overall quality was checked using agarose electrophoresis and a Qubit 2.0 fluorometer (Invitrogen, Carlsbad, CA, USA).

Twenty-four species per library were pooled in equimolar ratios (overall 18 libraries were prepared). In-solution sequence capture was done using MYbaits custom probes (Arbor Biosciences, formerly MYcroarray, Ann Arbor, MI, USA) following the manufacturer's protocol with a hybridization time of 26 hours, 12 cycles of PCR enrichment using KAPA HiFi HotStart DNA Polymerase (ThermoFisher Scientific, Wilmington, DE, USA), purification with the QIAquick PCR Purification kit (Qiagen, Venlo, Netherlands) and quantification using Qubit 2.0 (Invitrogen, Carlsbad, CA, USA).

*Processing of raw reads*. We used the HybPhyloMaker v. 1.6.4 pipeline [3] for raw read filtering, mapping of the filtered reads to a pseudoreference (exon sequences used for probe design, separated by strings of 400Ns) and construction of gene alignments. PhiX reads were removed with Bowtie 2 [4], SAMtools [5] and bam2fastq (https://gsl.hudsonalpha.org/ information/ software/bam2fastq). Adapter trimming and quality filtering was done using Trimmomatic [6], and FastUniq [7] was used for duplicate read removal. Mapping of filtered reads to a pseudoreference was performed using BWA [8]. A consensus sequence by majority rule consensus (majority threshold of 0.6) was generated with Kindel (https://github.com/bede/ kindel) and was then compared to the original exon sequences for each sample using BLAT [9] with a minimum sequence identity of 85. Consensus sequences were aligned using MAFFT 7 [10] with default settings and sequences from nuclear exons of the same gene assembly were concatenated into the particular genes using AMAS [11].

**Supplementary Figures**

**Fig. S1.** Flow-cytometric histograms and scatter plots for two species of the genus *Lepanthes* showing a different type of endoreplication.

**Fig. S2.** Ancestral state reconstructions of endoreplication type using stochastic character mapping.

**Fig. S3.** Ancestral state reconstructions of endoreplication type using a full BiSSE model.

**Fig. S4.** Ancestral state reconstructions of endoreplication type using a BiSSE model with equal extinction rates.

**Fig. S5.** Ancestral state reconstructions of endoreplication type using a BiSSE model with equal speciation rates.

**Fig. S6.** Credible intervals of differences from the posterior distribution of samples taken from BiSSE analyses.

**Fig. S7.** Species tree topologies obtained via the ASTRAL approach (a) and ML concatenation (b) of Pleurothallidinae.

**Fig. S8.** Divergence time estimates and 95% credibility intervals (CIs) inferred for all nodes of Pleurothallidinae chronograms.

**Fig. S9.** Mean phylorate plot of net diversification from BAMM analysis.

**Fig. S10.** The nine most common credible shift sets from BAMM analysis.

**Fig. S11.** Relative evidence of shifts in endoreplication mode inferred from the posterior distribution of BAMM results.

**Fig. S12.** Credible intervals of differences from the posterior distribution of samples taken from BAMM analyses and divided according to the type of endoreplication into partial (PE), conventional (CE) and mixed clades.

**Fig. S13.** Posterior distribution of state-dependent rates extracted from BiSSE analyses with equal extinction rates.

**Fig. S14.** State-dependent diversification rates extracted from BAMM analyses corresponding to three types of clades according to the harboured type or types of endoreplication – clades with exclusively conventional endoreplication (CE clades), clades with exclusively partial endoreplication (PE clades) and clades with both types (mixed clades).

**Supplementary tables**

**Table S1.** List of genera according to the adopted taxonomic concept supplemented with the number of species included, total species richness, endoreplication type, and data inferred from ASTRAL topology.

**Table S2.** List of specimens included in the study supplemented by the type of endoreplication, morphological traits (only for ingroup taxa) and HybSeq statistics (number of reads, number of genes and missing data). Coding for outgroup taxa starts with prefix "OUT", for ingroup with "P". (separate Excel table)

**Table S3.** Summary characteristics for clades corresponding to genera with significantly increased diversification rates.

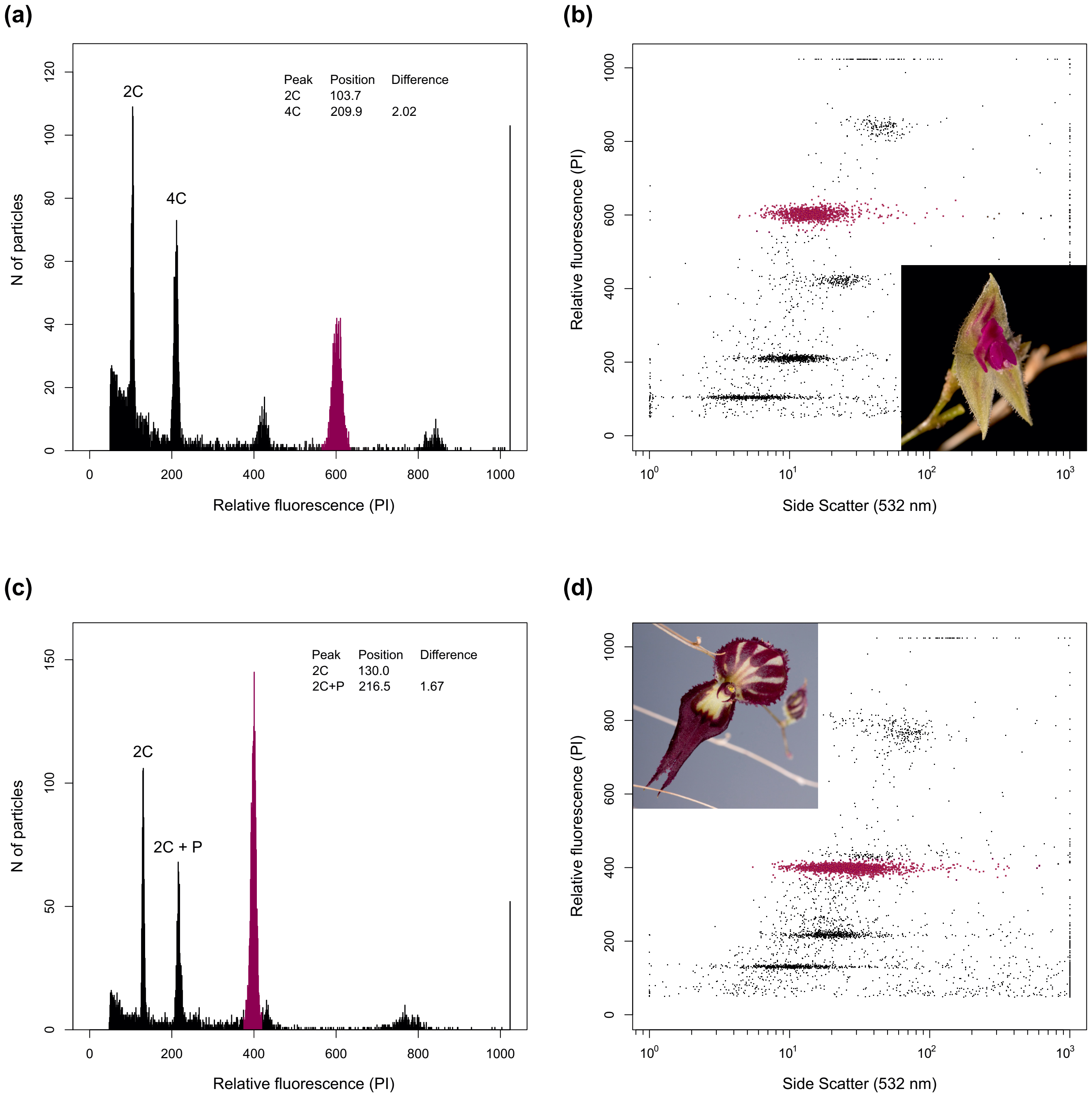


**Fig. S1.** Flow-cytometric histograms and scatter plots for two species of the genus *Lepanthes* showing a different type of endoreplication – *L. acarina* with conventional (a) and (b), and *L. katleri* with partial (c) and (d) endoreplication. The peak positions corresponding to subsequent fractions of nuclei of samples show the two-fold difference for (a) and (b), and 1.7-fold difference for (c) and (d). The fluorescence signal of the internal standard (*Solanum pseudocapsicum)* is purple. Labeling of peaks follows the common practice for both types of endoreplication.

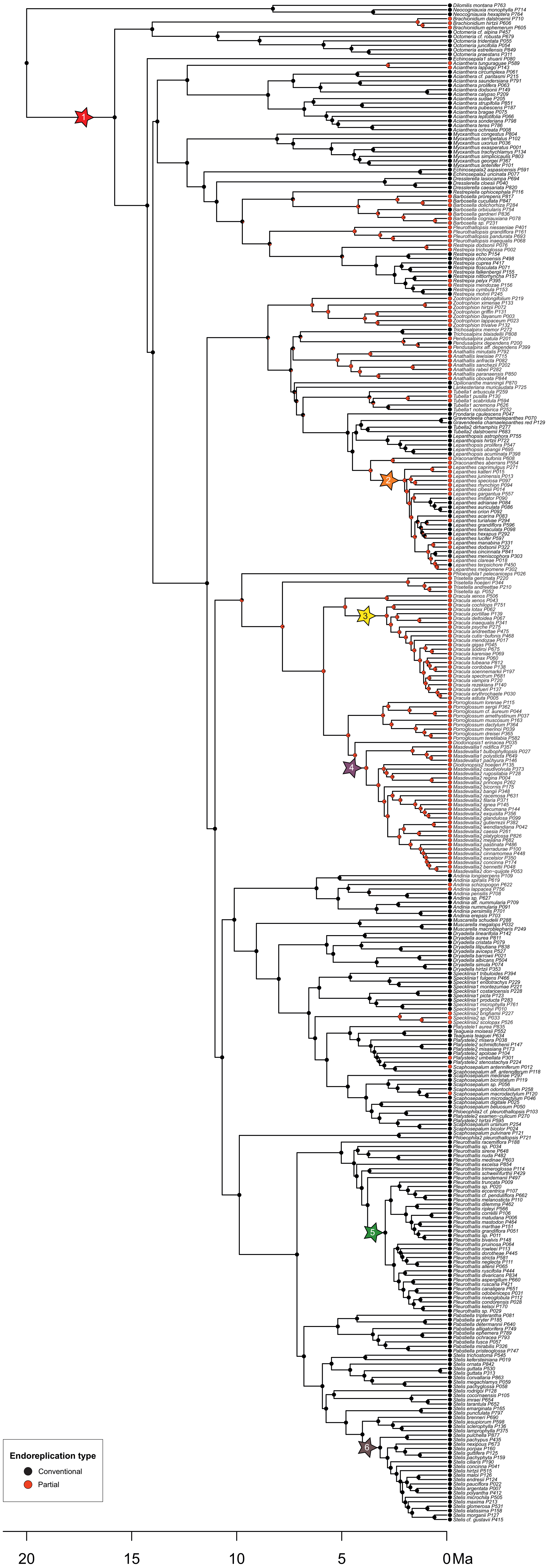


**Fig. S2.** Ancestral state reconstructions of endoreplication type using stochastic character mapping. Asterisks denote the position of the most credible shifts.

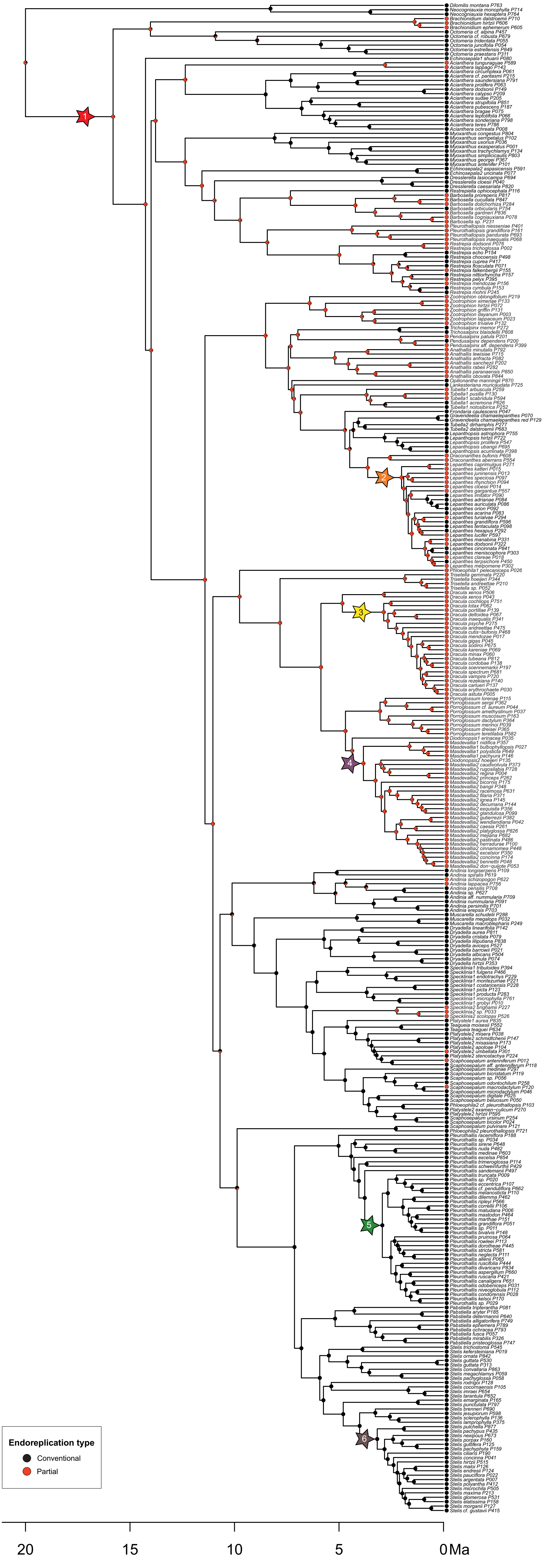


**Fig. S3.** Ancestral state reconstructions of endoreplication type using a full BiSSE model. Asterisks denote the position of the most credible shifts.

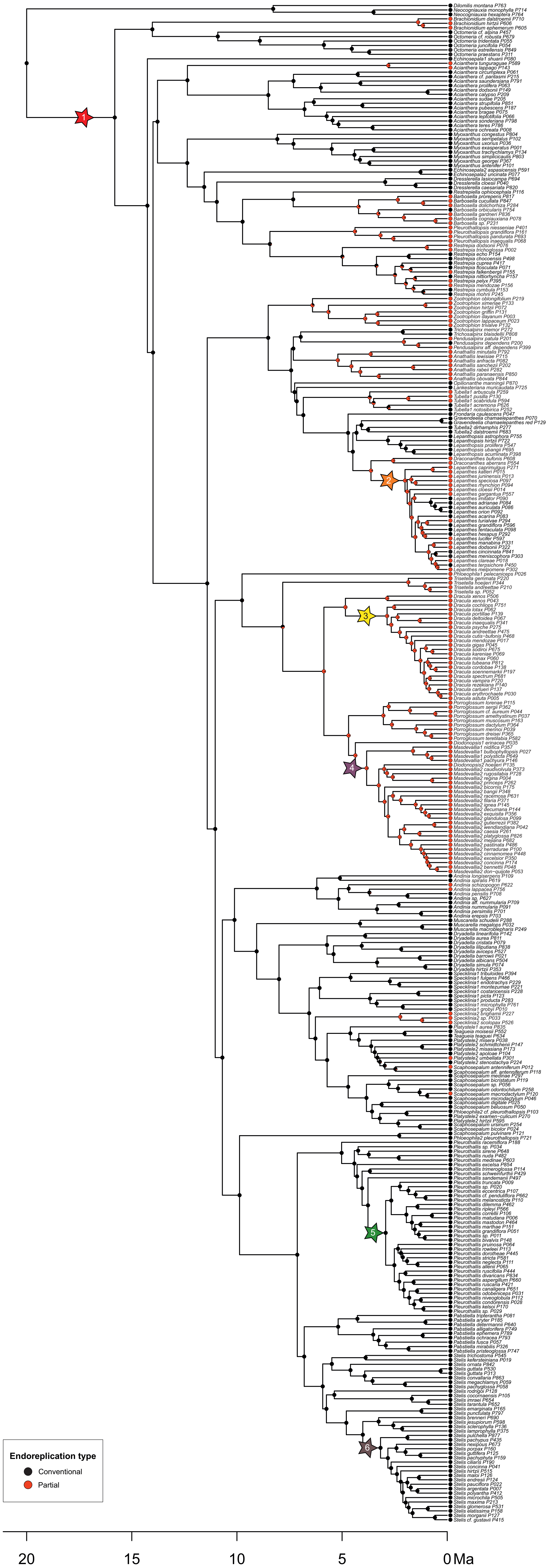


**Fig. S4.** Ancestral state reconstructions of endoreplication type using a BiSSE model with equal extinction rates. Asterisks denote the position of the most credible shifts.

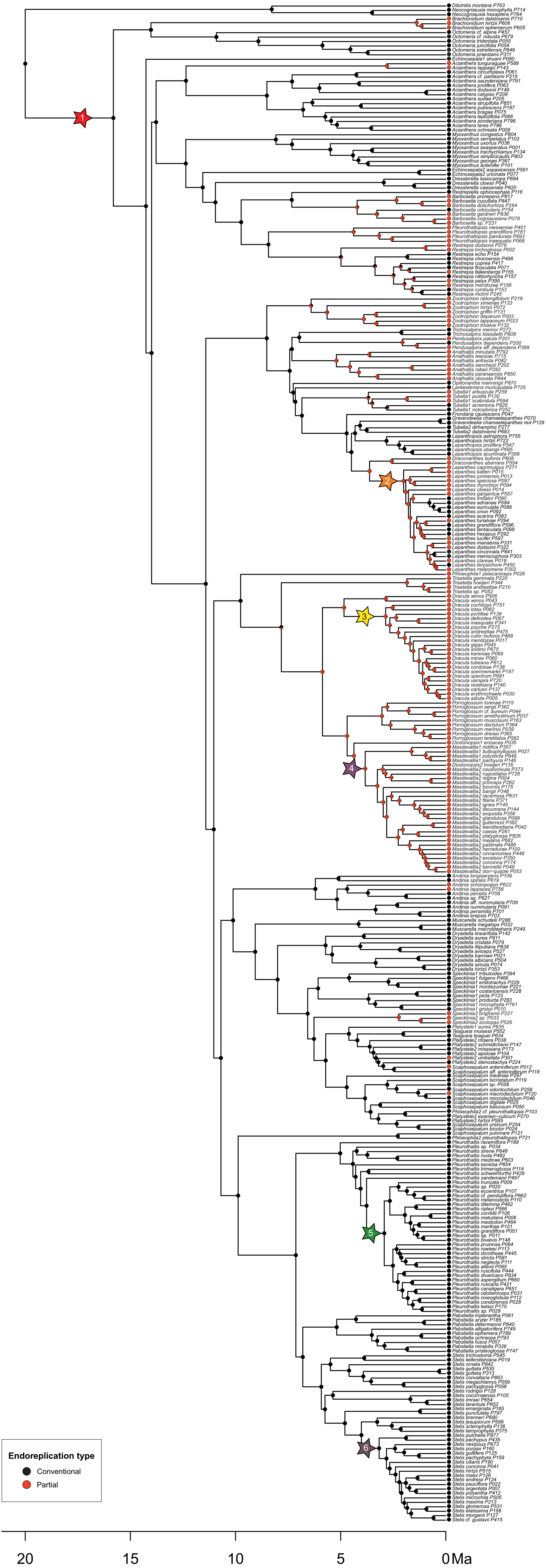


**Fig. S5.** Ancestral state reconstructions of endoreplication type using a BiSSE model with equal speciation rates. Asterisks denote the position of the most credible shifts.

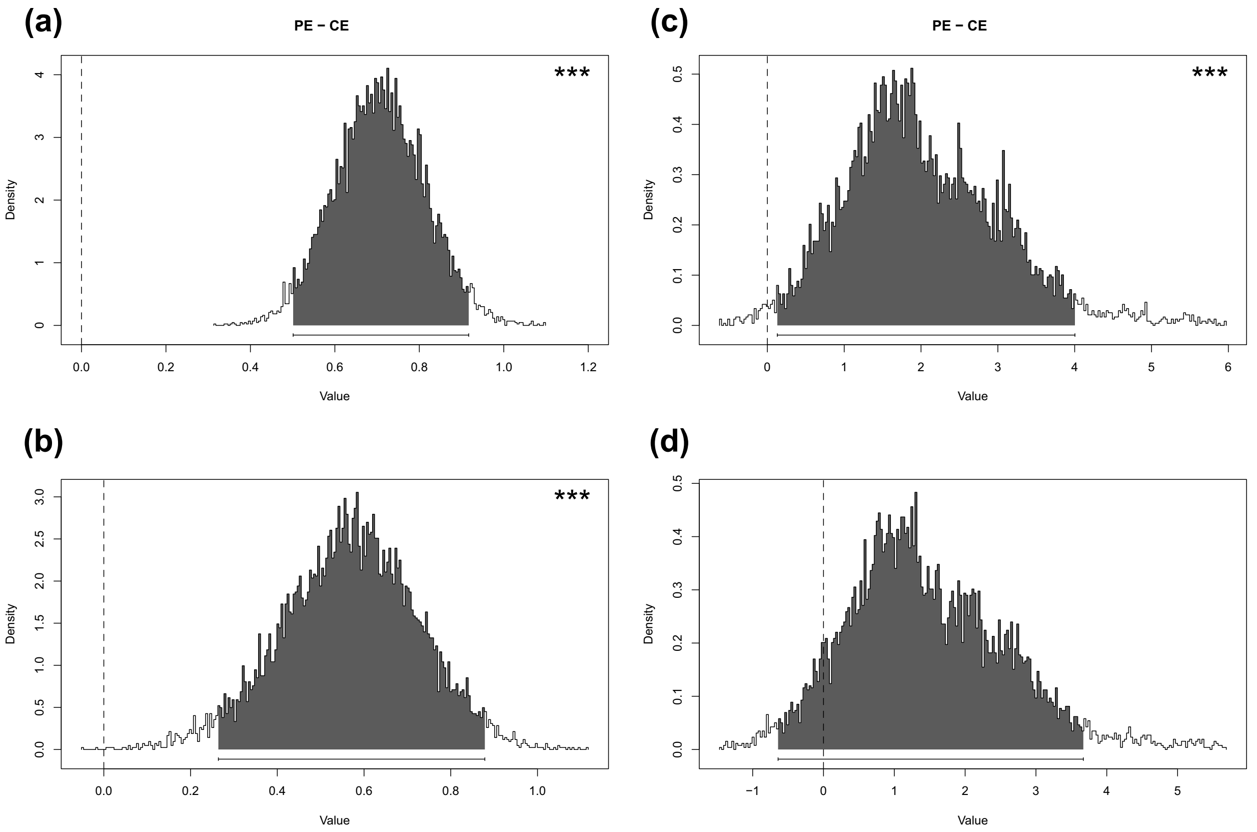


**Fig. S6.** Credible intervals of differences from the posterior distribution of samples taken from BiSSE analyses. The graphs compare the rates between the taxa with conventional (CE) and partial (PE) endoreplication for (a) a model with equal extinction rates, where differences in speciation and net diversification rates are the same, and a full model with differences in (b) net diversification, (c) speciation and (d) extinction rates. The dotted black line indicates a zero difference between samples. Black-shaded regions represent probability intervals. Significance is calculated as the percentage of credible differences that do not overlap with zero, represented as * ≥ 95%, ** ≥ 99% and *** ≥ 99.9%; one-tailed.

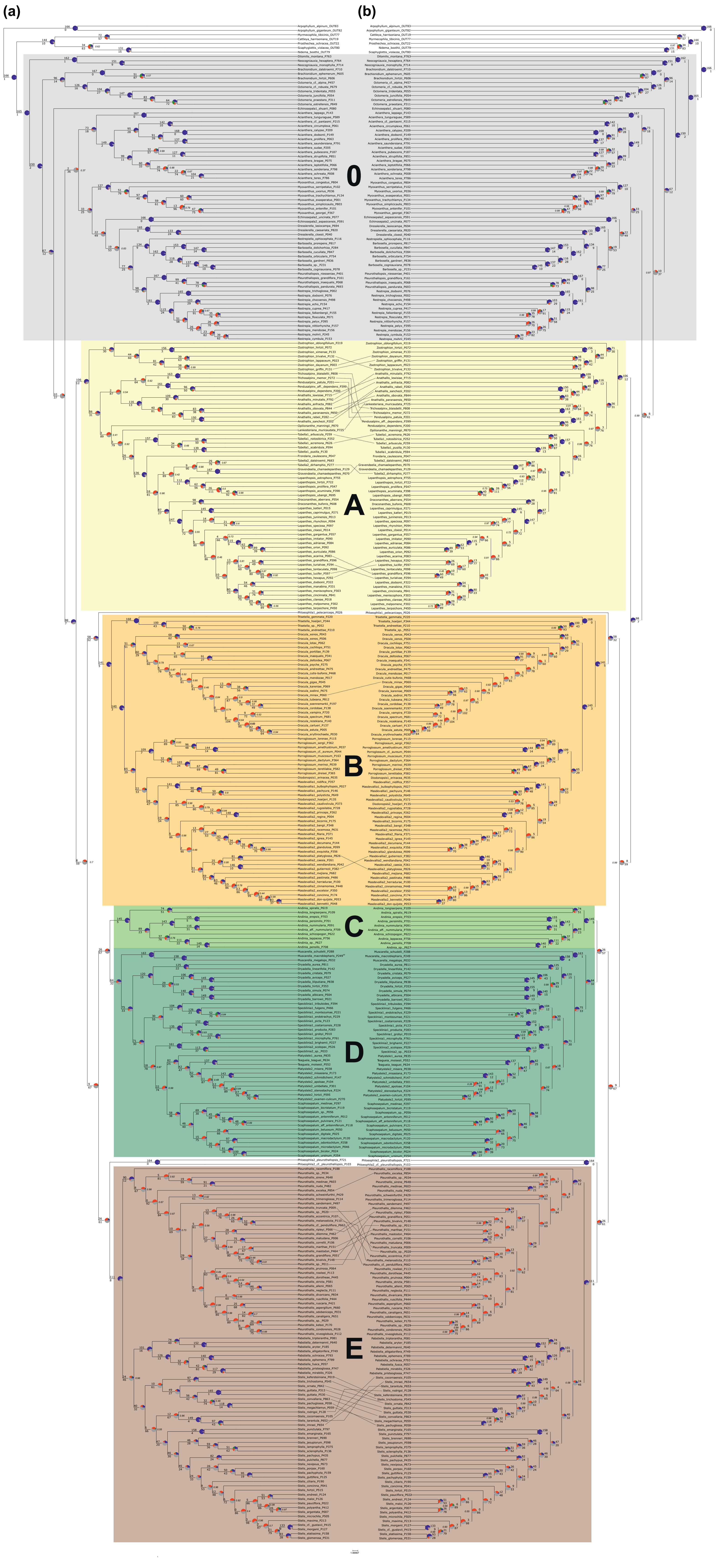


**Fig. S7.** Species tree topologies obtained via the ASTRAL approach (a) and ML concatenation (b) of Pleurothallidinae. Both trees are supplemented with a summary of gene trees that are in conflict with or are concordant with species tree topology. For each branch, the top number indicates the number of gene trees concordant with the species tree at that node, and the bottom number indicates the number of gene trees in conflict with that clade in the species tree. The pie charts at each node present the proportion of gene trees that support that clade (blue), the proportion that support the main alternative for that clade (green), the proportion that support the remaining alternatives (red), and the proportion that inform (conflict or support) this clade that have less than 50% bootstrap support (grey). Bayesian posterior probabilities (PP) for the ASTRAL tree and SH-aLRT (SH) values for the ML concatenated tree are indicated in italics only for branches with support lower than 1 PP or 1 SH. The topological differences between two species trees are highlighted using a tanglegram, where matching accessions in the two trees are joined by an edge.

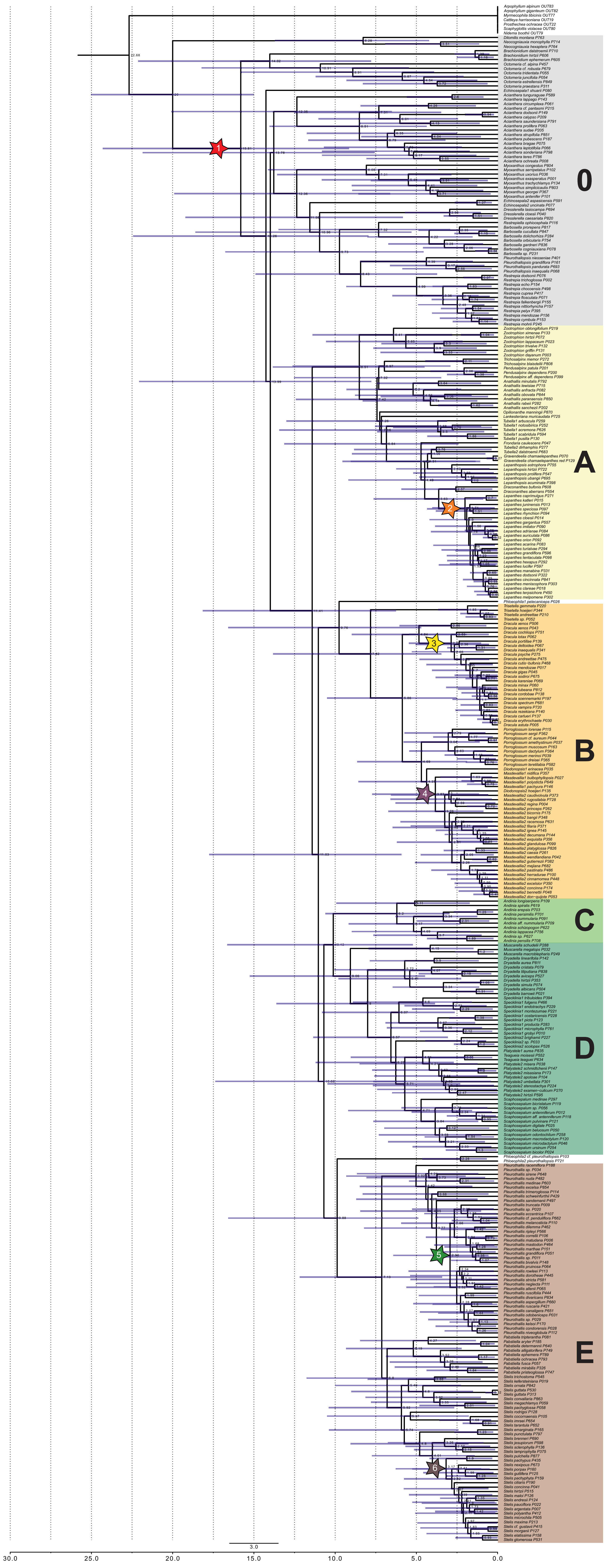


**Fig. S8.** Divergence time estimates and 95% credibility intervals (CIs) inferred for all nodes of Pleurothallidinae chronograms. Asterisks denote the position of the most credible shifts.

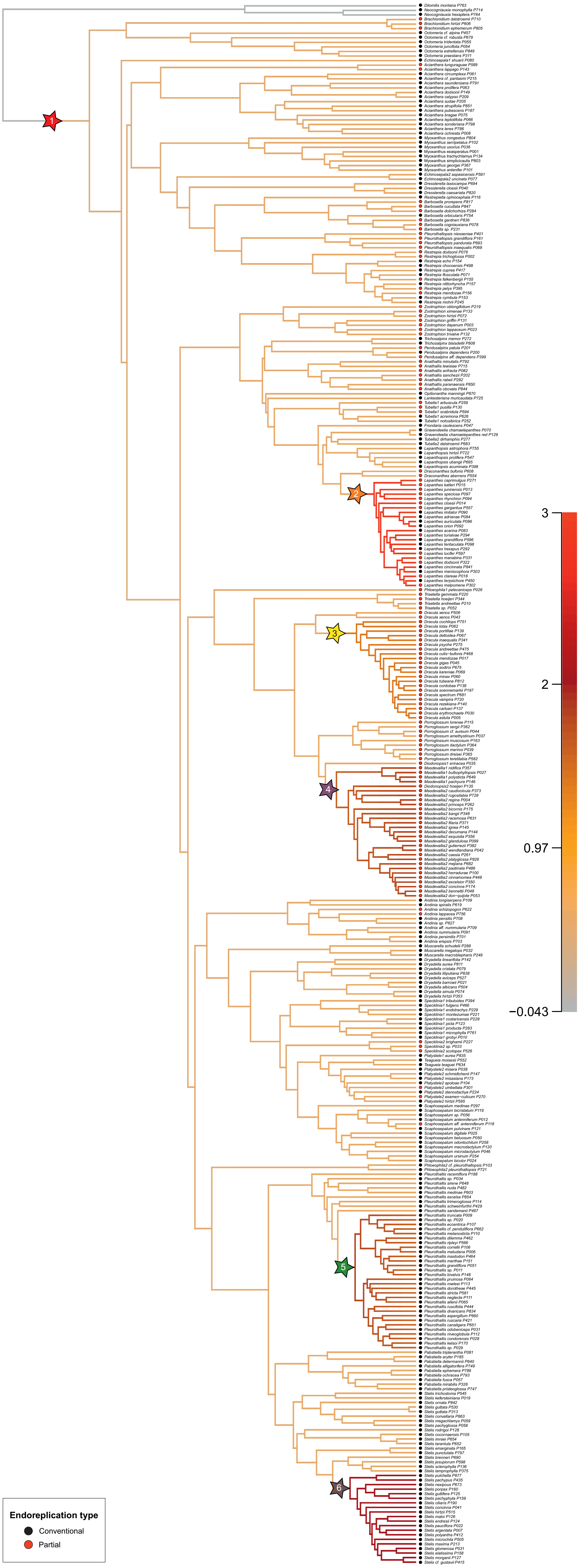


**Fig. S9.** Mean phylorate plot of net diversification from BAMM analysis. Asterisks denote the position of the most credible shifts, red circles denote partial endoreplication and black circles denote conventional endoreplication.

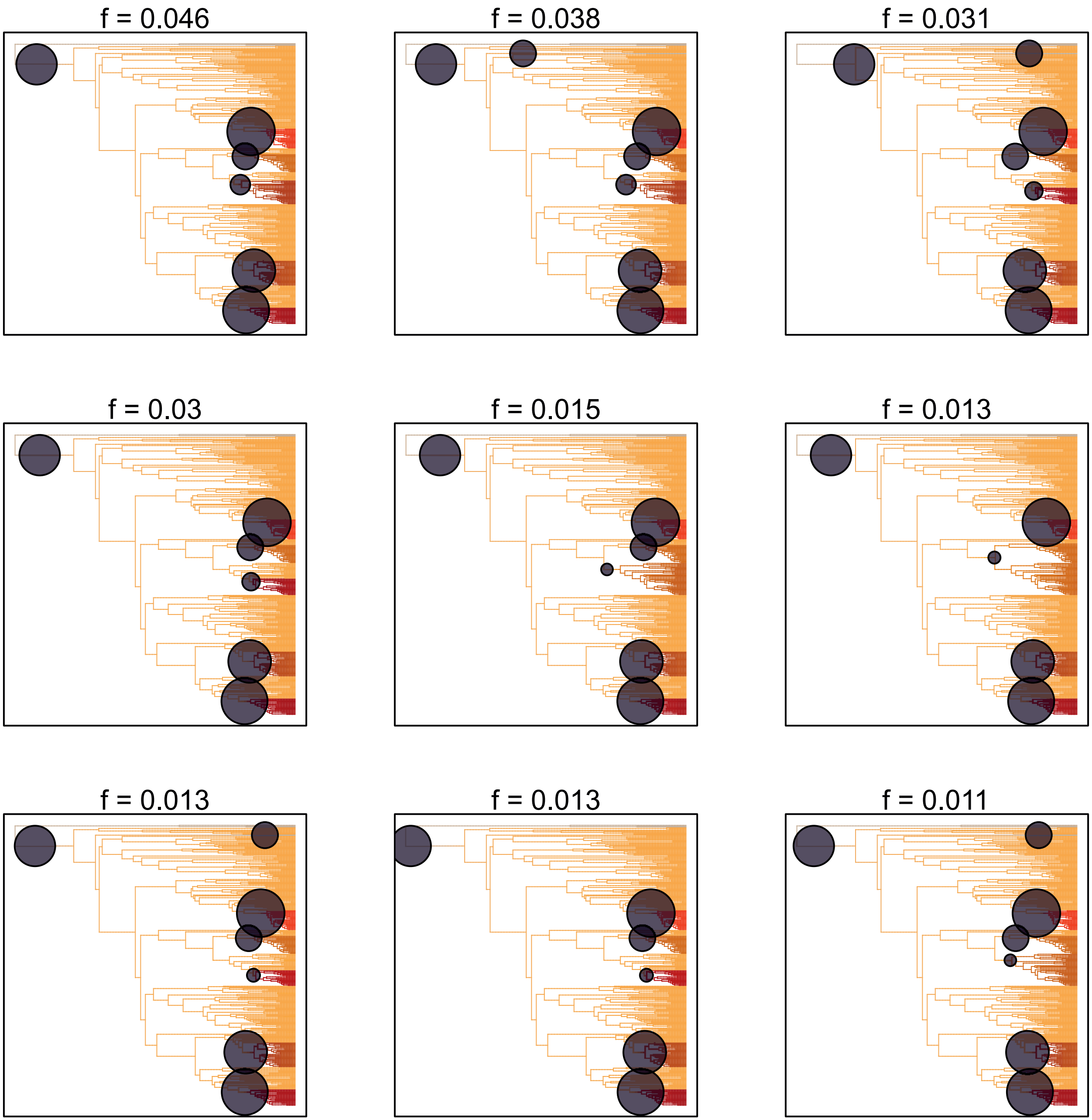


**Fig. S10.** The nine most common credible shift sets from BAMM analysis.

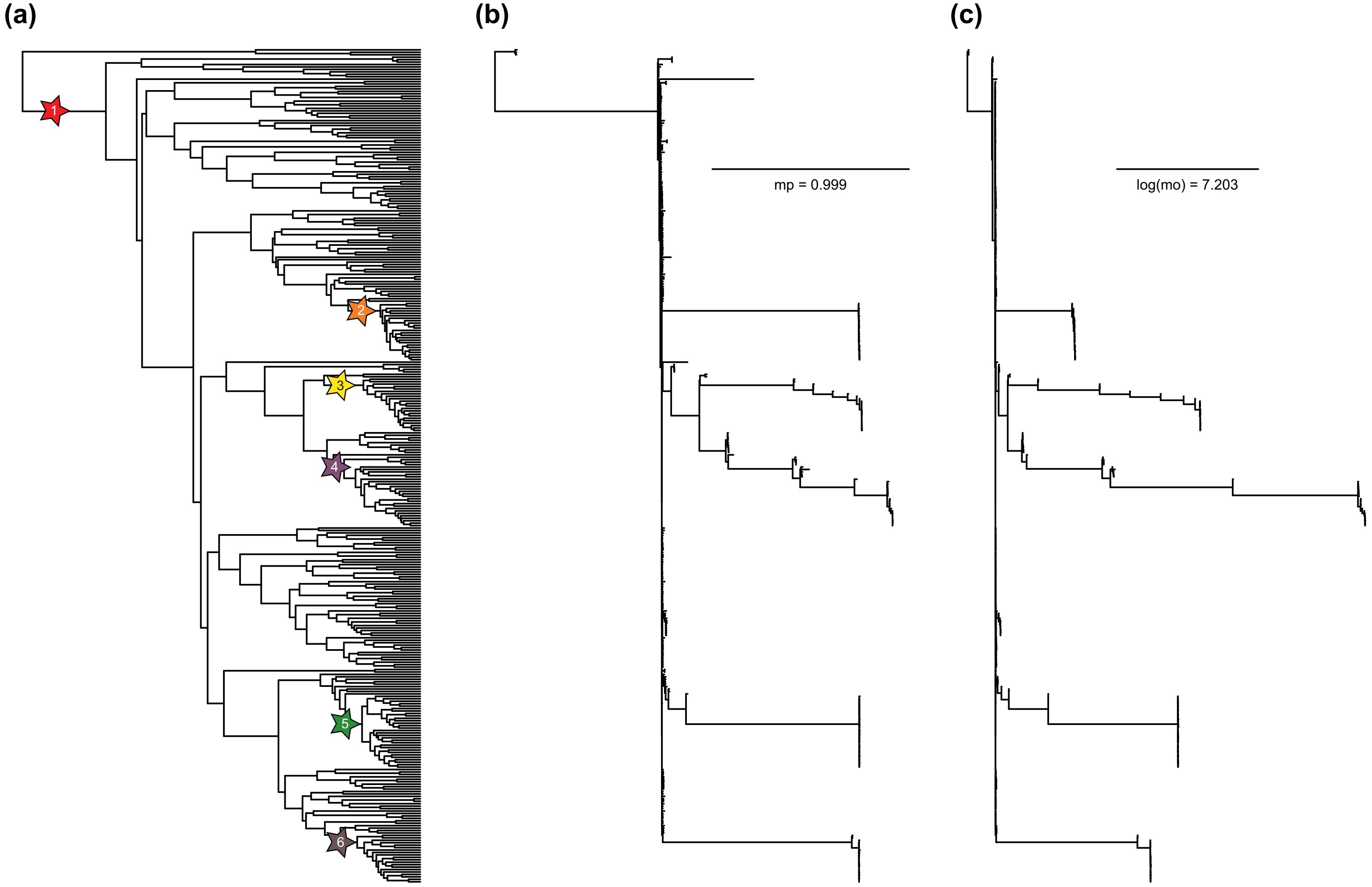


**Fig. S11.** Relative evidence of shifts in endoreplication mode inferred from the posterior distribution of BAMM results. (a) Original tree with asterisks denoting the position of the most credible shifts, (b) marginal probability tree with branch length corresponding to the probability that a shift occurred along the branch, (c) marginal odds tree with branch lengths corresponding to branch-specific pseudo-Bayes factors.

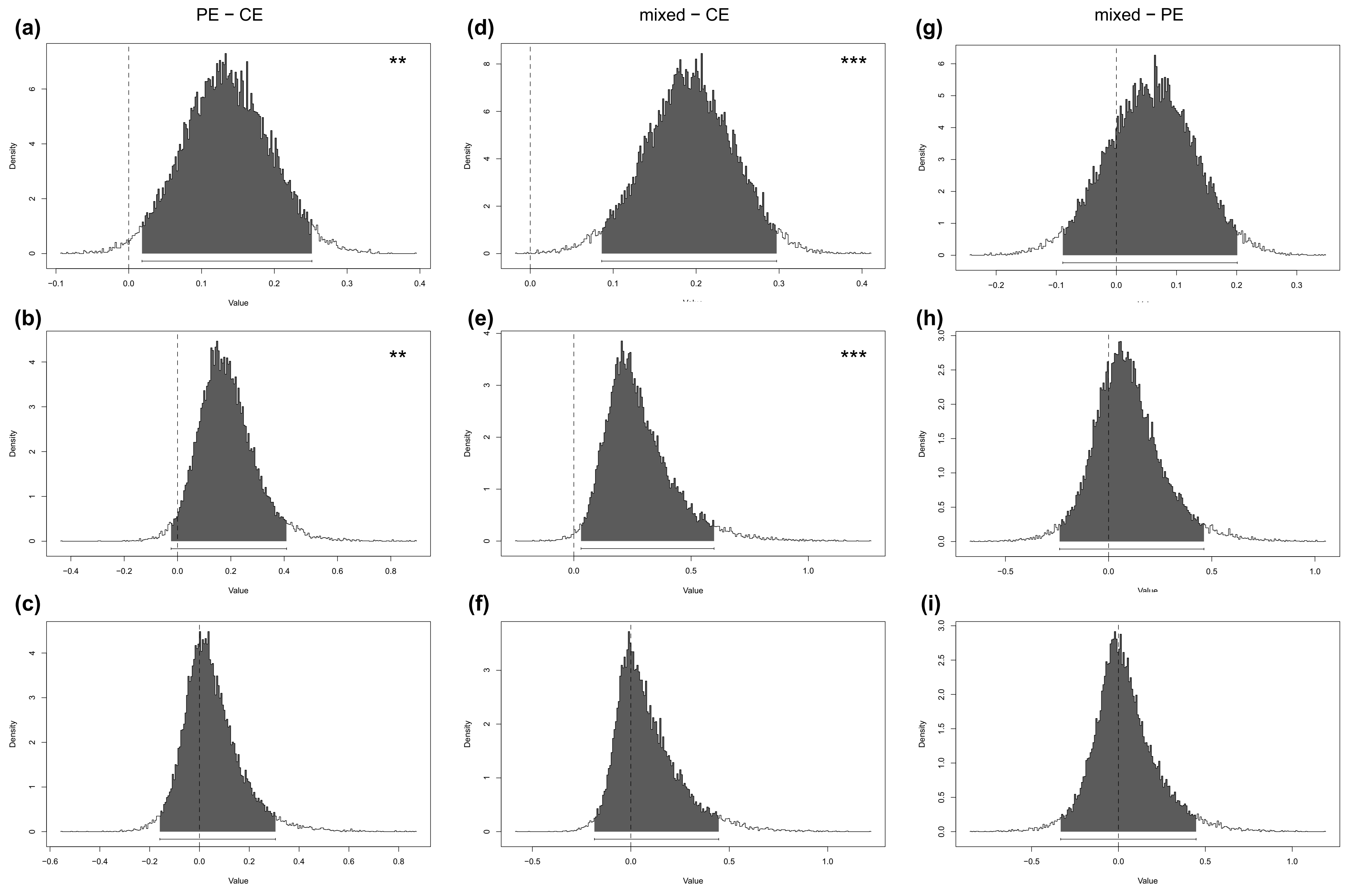


**Fig. S12.** Credible intervals of differences from the posterior distribution of samples taken from BAMM analyses and divided according to the type of endoreplication into partial (PE), conventional (CE) and mixed clades. Graphs in the left column compare PE and CE clades for (a) net diversification rates, (b) speciation rates, and (c) extinction rates. Graphs in the middle column compare mixed and CE clades for (d) net diversification rates, (e) speciation rates, and (f) extinction rates. Graphs in the right column compare (g) net diversification rates, (h) speciation rates, and (i) extinction rates for mixed and PE clades. The dotted black line indicates a zero difference between samples. Black-shaded regions represent probability intervals. Significance is calculated as the percentage of credible differences that do not overlap with zero, represented as * ≥ 95%, ** ≥ 99% and *** ≥ 99.9%; one-tailed.

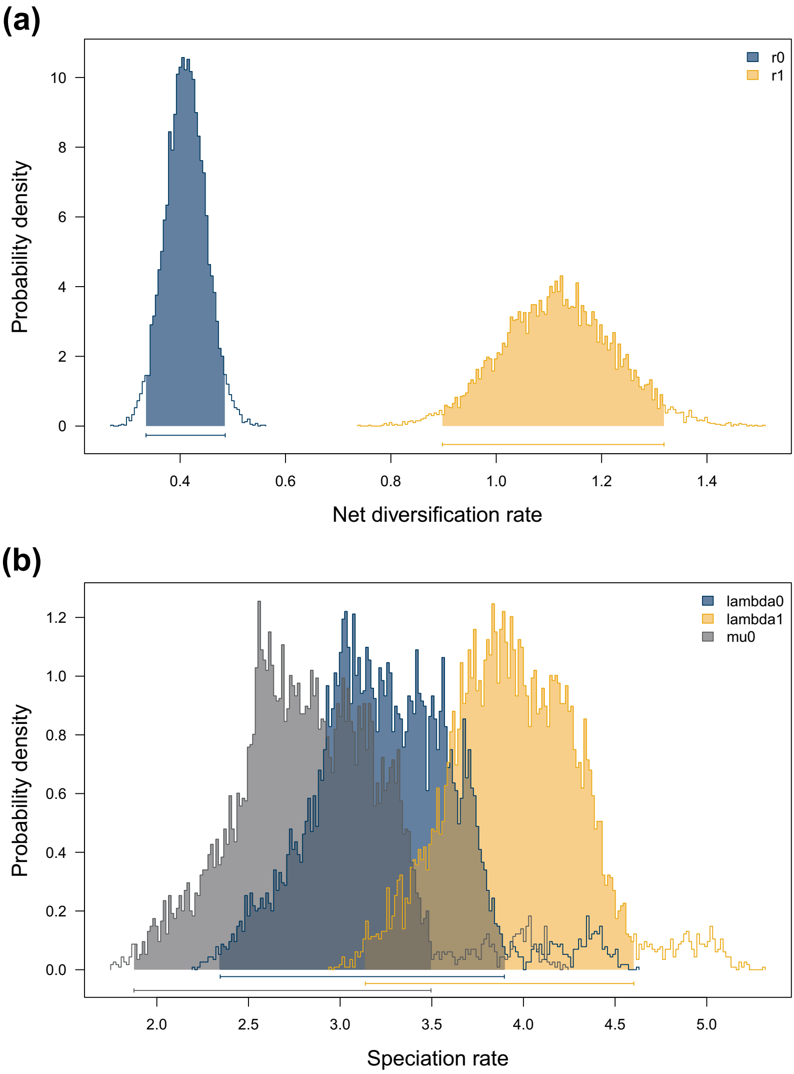


**Fig. S13.** Posterior distribution of state-dependent rates extracted from BiSSE analyses with equal extinction rates. The graphs show the posterior distribution of rates for (a) net diversification and (b) speciation with extinction, coloured by character state (type of endoreplication) or by diversification or extinction rates, respectively.

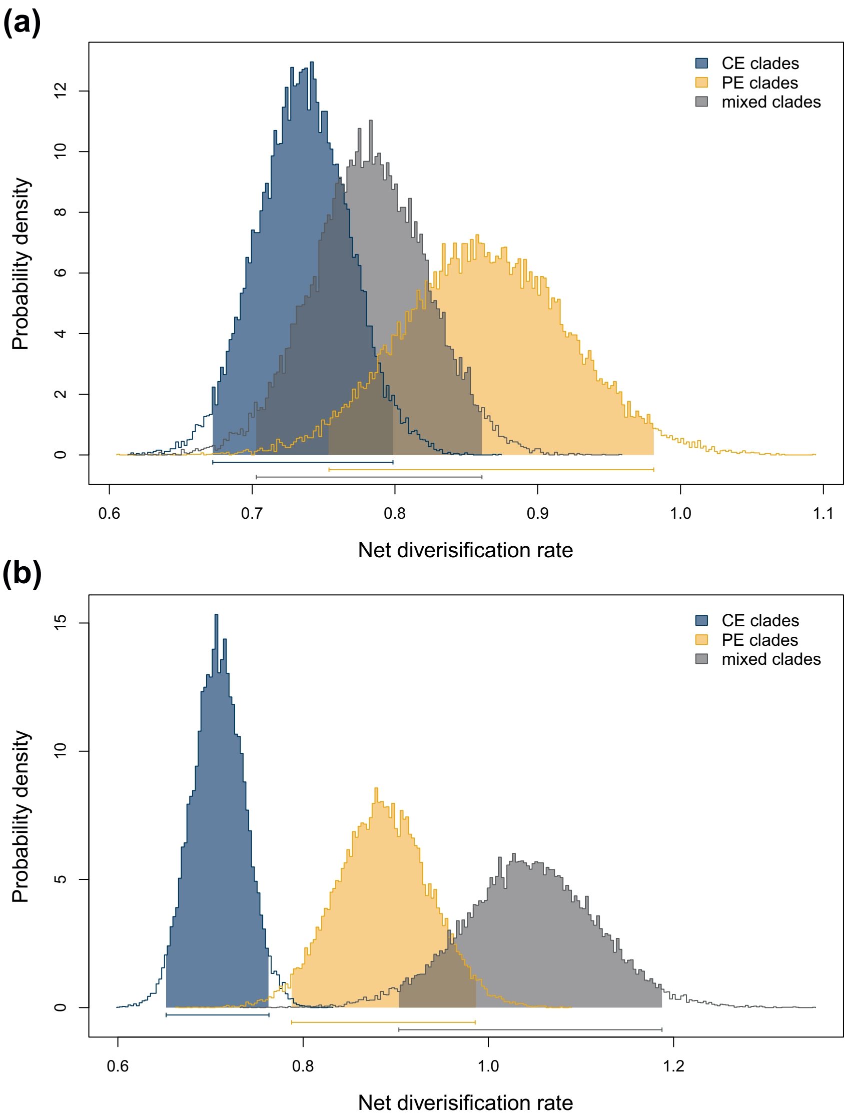


**Fig. S14.** State-dependent diversification rates extracted from BAMM analyses corresponding to three types of clades according to the harboured type or types of endoreplication – clades with exclusively conventional endoreplication (CE clades), clades with exclusively partial endoreplication (PE clades) and clades with both types (mixed clades). The graphs show the posterior distribution of net diversification rates for (a) a rough estimation of clades according to the generic concept and (b) the finest possible determination of mixed clades; coloured by the character state of clades.

**Table S1.** List of genera according to the adopted taxonomic concept supplemented with the number of species included, total species richness, endoreplication type (PE – partial endoreplication, CE – conventional endoreplication, mixed – both types), support for genera in ASTRAL species tree in terms of the number gene trees that are in concordance or in conflict with the species tree topology and their placement in six groups introduced in Fig. S2.

| Genus | Species analysed | Total number of recognized species | Endoreplication type | Number of gene trees in concordance/conflict with the monophyly of the group | Group |
|---|---|---|---|---|---|
| *Acianthera* | 16 | 292 | mixed | 73/18 | 0 |
| *Anathallis* | 7 | 147 | PE | 116/13 | A |
| *Andinia* | 10 | 74 | mixed | 145/5 | C |
| *Atopoglossum* | NA | 3 | NA | NA | NA |
| *Barbosella* | 7 | 19 | Mixed | 162/2 | 0 |
| *Brachionidium* | 3 | 82 | PE | 167/0 | 0 |
| *Chamelophyton* | NA | 1 | NA | NA | NA |
| *Dilomilis* | 1 | 5 | CE | 162/1 | 0 |
| *Diodonopsis* | 2 | 6 | PE | *Diodonopsis*1 22/53 | B |
| | | | | *Diodonopsis*2 5/85 | B |
| *Draconanthes* | 2 | 2 | PE | 98/28 | A |
| *Dracula* | 23 | 141 | PE | 51/51 | B |
| *Dresslerella* | 3 | 14 | CE | 165/0 | 0 |
| *Dryadella* | 9 | 58 | CE | 120/15 | D |
| *Echinosepala* | 3 | 14 | CE | *Echinosepala*1 67/12 | 0 |
| | | | | *Echinosepala*2 166/0 | 0 |
| *Frondaria* | 1 | 1 | CE | 136/5 | A |
| *Gravendeelia* | 1 | 1 | CE | 167/0 | A |
| *Lankesteriana* | 1 | 24 | CE | 3/79 | A |
| *Lepanthes* | 24 | 1134 | mixed | 147/1 | A |
| *Lepanthopsis* | 5 | 47 | CE | 112/11 | A |
| *Masdevallia* | 28 | 652 | PE | *Masdevallia*1 141/3 | B |
| | | | | *Masdevallia*2 19/54 | B |
| *Muscarella* | 3 | 52 | CE | 163/1 | D |
| *Myoxanthus* | 8 | 49 | CE | 127/10 | 0 |
| *Neocogniauxia* | 2 | 2 | CE | 151/9 | 0 |
| *Octomeria* | 6 | 165 | CE | 126/16 | 0 |
| *Opilionanthe* | 1 | 1 | CE | 3/89 | A |
| *Pabstiella* | 9 | 128 | CE | 101/16 | E |
| *Pendusalpinx* | 3 | 7 | mixed | 161/0 | A |
| *Phloeophila* | 3 | 11 | mixed | *Phloeophila*1 58/32 | / |
| | | | | *Phloeophila*2 164/0 | / |
| *Platystele* | 9 | 116 | mixed | *Platystele*1 82/25 | D |

| | | | | | |
|---|---|---|---|---|---|
| | | | | *Platystele*2 52/36 | D |
| *Pleurothallis* | 39 | 541 | CE | 90/12 | E |
| *Pleurothallopsis* | 4 | 19 | PE | 167/0 | 0 |
| *Porroglossum* | 9 | 54 | PE | 109/7 | B |
| *Pseudolepanthes* | NA | 10 | NA | NA | NA |
| *Restrepia* | 12 | 61 | mixed | 161/3 | 0 |
| *Restrepiella* | 1 | 4 | CE | 134/9 | 0 |
| *Sansonia* | NA | 2 | NA | NA | NA |
| *Scaphosepalum* | 13 | 54 | mixed | 58/38 | D |
| *Specklinia* | 12 | 103 | mixed | *Specklinia*1 30/55 | D |
| | | | | *Specklinia*2 161/0 | D |
| *Stelis* | 38 | 1253 | CE | 40/32 | E |
| *Stellamaris* | NA | 1 | NA | NA | NA |
| *Teagueia* | 2 | 18 | CE | 137/7 | D |
| *Tomzanonia* | NA | 1 | NA | NA | NA |
| *Trichosalpinx* | 2 | 29 | CE | 163/0 | A |
| *Trisetella* | 4 | 26 | PE | 168/0 | B |
| *Tubella* | 7 | 73 | mixed | *Tubella*1 157/0 | A |
| | | | | *Tubella*2 29/79 | A |
| *Zootrophion* | 7 | 28 | PE | 71/30 | A |
| TOTAL | 340 | 5525 | | | |

**Table S3.** Summary characteristics for clades corresponding to genera with significantly increased diversification rates. The table is supplemented with information on the number of gene trees that are in concordance or conflicting with the monophyly of the clades and mean divergence times with confidence intervals (CI). Clades within the genus with increased diversification rates are labeled as *DIV.

| Group | Number of gene trees in concordance/conflict with the monophyly of the group | Divergence time [mya] | 95% CI [mya] |
|---|---|---|---|
| *Lepanthes* | 147/1 | 2.4 | 0.4–4.3 |
| *Dracula* (excl. *D. xenos*) | 60/29 | 2.8 | 1–5.4 |
| *Masdevalia* 1 & 2 (incl. *Diodonopsis* 2) | 28/47 | 3.8 | 1–7.6 |
| *Pleurothalis* | 90/12 | 5 | 1.9–9.3 |
| *Pleurothalis* DIV* | 29/34 | 2.9 | 0.5–5.4 |
| *Stelis* | 40/32 | 5.9 | 2.6–10.3 |
| *Stelis* DIV* | 45/24 | 3.1 | 1–6 |
